# A Regression Framework for Brain Network Distance Metrics

**DOI:** 10.1101/2021.02.26.432910

**Authors:** Chal E. Tomlinson, Paul J. Laurienti, Robert G. Lyday, Sean L. Simpson

## Abstract

Analyzing brain networks has long been a prominent research topic in neuroimaging. However, statistical methods to detect differences between these networks and relate them to phenotypic traits are still sorely needed. Our previous work developed a novel permutation testing framework to detect differences between two groups. Here we advance that work to allow both assessing differences by continuous phenotypes and controlling for confounding variables. To achieve this, we propose an innovative regression framework to relate distances between brain network features to functions of absolute differences in continuous covariates and indicators of difference for categorical variables. We explore several similarity metrics for comparing distances between connection matrices, and adapt several standard methods for estimation and inference within our framework: Standard F-test, F-test with individual level effects (ILE), Feasible Generalized Least Squares (FGLS), and Permutation. Via simulation studies, we assess all approaches for estimation and inference while comparing them with existing Multivariate Distance Matrix Regression (MDMR) methods. We then illustrate the utility of our framework by analyzing the relationship between fluid intelligence and brain network distances in Human Connectome Project (HCP) data.

**Highlights:**

- Related distances between connection matrices to differences in covariates.
- Adapted methods for estimation and inference in this framework.
- Assessment of methods and distance metrics via simulation.
- Compared our methods to existing MDMR methods via simulation.
- Analysis of the HCP data with the best approach for each distance metric.

## 1. Introduction

As brain network analyses have become popular in recent years, neuroimaging researchers often face the need to statistically compare brain networks (Simpson et al., 2013a). Many approaches for relating brain networks to clinical outcomes or demographical variables have been developed. Such methods include, but are not limited to traditional network models (e.g., exponential random graph models (Lehmann et al., 2021; Simpson et al., 2012, 2011)), tensor regression works on brain network (e.g. (Zhang et al., 2018, 2019)), Bayesian approaches (e.g. (Dai and Guo, 2017; Wang et al., 2017)), and statistical learning techniques (RC et al., 2015; Varoquaux and Craddock, 2013; Xia et al., 2020). Despite the advances made, analysis methods are still needed that enable the comparison of networks while incorporating topological features inherent in each individual’s network. In order to develop such an analysis, we can exploit the fact that brain networks often exhibit consistent organizations across subjects. For example, a number of studies have reported that nodes with particular characteristics (e.g., high degree) tend to coincide at the same spatial locations across subjects (Hagmann et al., 2008; Hayasaka and Laurienti, 2010; Moussa et al., 2011; van den Heuvel et al., 2008). Although the set of such nodes may not be exactly the same across subjects, there are large areas of overlap (Hagmann et al., 2008; Hayasaka and Laurienti, 2010). Furthermore, our study on network modules, or communities of highly-interconnected nodes, indicated that some building blocks of resting-state functional brain networks exhibited remarkable consistency across subjects (Moussa et al., 2012). It has also been shown that such consistent organizations differ under different cognitive states (Deuker et al., 2009; Moussa et al., 2011; Rzucidlo et al., 2013) or in different groups of subjects (Bassett and Bullmore, 2009; Burdette et al., 2010; Liu et al., 2008; Meunier et al., 2009a; Rombouts et al., 2005; Stam et al., 2007; Yuan et al., 2010). Thus, an analysis method sensitive to such differences in spatial locations or patterns can assess network differences across the entire brain (as opposed to univariate edge-by-edge or node-by-node comparisons). Toward this end, in previous work we developed a permutation testing framework that detects whether the spatial location of network features (such as the location of high degree nodes) mapped back into brain space differs between two groups of networks, and whether distributions of topological properties vary by group (Simpson et al., 2013b). Despite the utility of this method, it has two major weaknesses. First, it cannot account for confounding variables. This means that while we can compare maps of hub regions, for example, between two populations, we cannot control for differences in characteristics such as age or education. Second, the method relies on dichotomous grouping to make comparisons. When making comparisons between groups with clear divisions (male vs. female), this is not an issue. However, it is impossible to assess if there is a significant relationship between network hub location and continuous measures, such as intelligence quotient (IQ) score or age.

To address these limitations, we propose an innovative regression framework to relate distances between brain network features to functions of absolute differences in continuous covariates and indicators of difference for categorical variables. We will consider several different types of metrics for establishing distances (i.e., similarity/dissimilarity) between networks. The first type compares degree distributions. We accomplish this by summarizing similarities in nodal cumulative degree distributions across multiple networks with the Kolmogorov-Smirnov statistic (K-S statistic), a measure that quantifies the distance between two cumulative distribution functions (Kolmogorov, 1933; Smirnov, 1948). The second type takes into account consistency of key node sets. We do so by summarizing similarities in node sets across multiple networks with the Jaccard index, a metric that quantifies similarity in partitions of a set (Joyce et al., 2010; Meunier et al., 2009b). The third type of metric measures similarity of nodal degree by employing Minkowski or Canberra distances between nodal degree vectors (Lance and Williams, 1966).

Within our regression framework we adapt several methods for estimation and inference: Standard F-test, F-test with individual level effects (ILE), Feasible Generalized Least Squares (FGLS), and Permutation. Each observation in the regression framework includes a “distance” between two individuals, so observations that share individuals are correlated. Thus, the standard F-test is generally not appropriate, but presented for comparison. The other methods attempt to deal with this correlation: 1) F-test with ILE—including fixed individual level effects within the regression, possibly rendering the F-test valid; 2) Mixed Model—including random individual level effects; 3) FGLS—proposing an artful way to estimate the covariance matrix (Aitken, 1936); and 4) Permutation—similarity and distance metrics have unknown distributions; thus, a permutation test may be most appropriate (Simpson et al., 2013b). Permutation tests for these models require permuting the residuals, and we will adapt recent methods to implement this approach (Kherad-Pajouh and Renaud, 2014, 2010).

In this paper, we detail our proposed regression framework, and discuss several methods for estimation and inference to be used on a variety of network similarity/dissimilarity metrics. We assess all combinations of methods and metrics within this framework using simulated fMRI data with known differences in connectivity matrix distributions. We then apply our framework to functional brain networks derived from the HCP dataset to investigate the relationship between fluid intelligence and network distances after accounting for known confounders.

## 2. Methods

Please note the following notational choices: bold font is used to denote vectors or matrices, *d* = number of nodes, 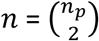 = number of observations, *n*_*p*_ = number of participants, *p* = number of covariates (including intercept, if included).

### 2.1. Step 1: Network Construction

Assuming fMRI connection matrices have already been obtained (see Figure 1 recreated from (Fornito et al., 2012; Simpson et al., 2013a)), let ***C***_***i***_ represent a weighted *d* × *d* connection matrix for individual *i*, with matrix entries ranging from 0 (indicating no connection between the respective nodes) to 1 (strongest connection). We only considered undirected networks, so matrices were symmetric, with the *# of row* = *# of columns* = *# of nodes* (methods are easily adaptable if directed networks are desired).

**Figure 1:**
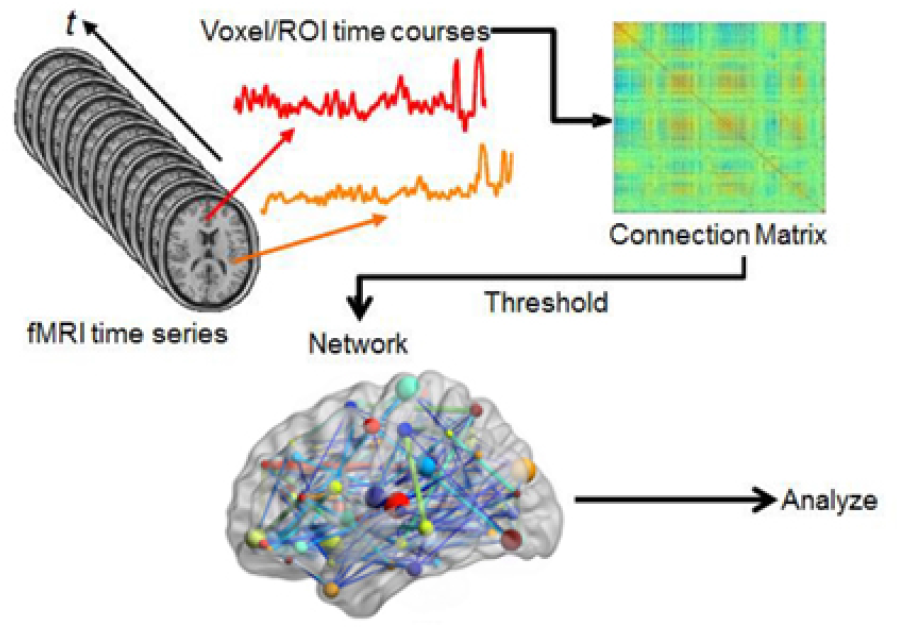
Schematic for generating brain networks from fMRI time series data—recreated from *(Fornito et al., 2012; Simpson et al., 2013a).* Functional connectivity between brain areas is estimated based on time series pairs to produce a connection matrix. A threshold is commonly applied to the matrix to remove negative and/or “weak” connections.

Let ***b***_***i***_ represent an *d* × 1 binary key node vector for individual *i*, with an entry of 1 representing a key node and 0 a node that is not key. As stated in our previous work (Simpson et al., 2013b), key nodes can be identified based on nodal characteristics such as high degree, high centrality, or other desired characteristics. Since key nodes were compared across subjects, it was important to employ the same criterion in all of the networks (e.g., top 10% highest centrality, node degree >200, etc.). Alternatively, key nodes can be identified as those belonging to a particular module or network community, a collection of highly interconnected nodes. The resulting key nodes would form a set and the consistency of the spatial location of the nodes could be compared across groups. An example visualization of key node sets from voxel-based networks in brain space is given in Figure 2--recreated from (Simpson et al., 2013b).

**Figure 2:**
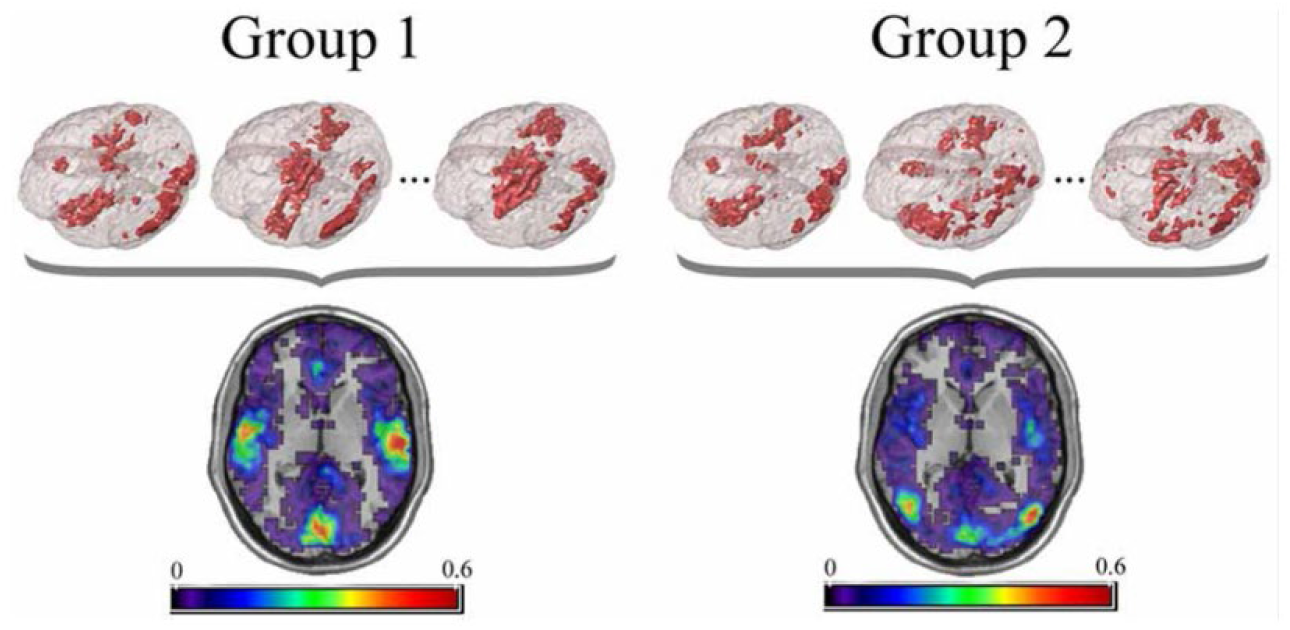
Example visualization of key node sets from voxel-based networks in brain space. The 3D brain images (Top) are 3 representative subjects from each group with the key node sets defined to be those with the top 20% highest degree. Overlap maps (Bottom) show the proportion of subjects with key nodes in given areas. Figure recreated from *(Simpson et al., 2013b).*

We constructed an *d* × 1 weighted nodal degree vector ***d***_***i***_ by summing ***C***_***i***_ over its rows (or equivalently, columns). Each entry in vector ***d***_***i***_ represented the “degree” of its respective node for individual *i*. It should be noted here that when the binary key node vector has to do with node degree, say 20% highest degree as will used later, ***b***_***i***_ is just a binarized version of ***d***_***i***_.

As was described for the binary key node vector in the last paragraph, other weighted nodal vectors could have been used. For simplicity, we focused on degree within the methods and simulation sections. It should also be noted that none of the methods employed here are specific to nodal vectors. That is, these methods could also be implemented on differences between connection matrices, for example.

### 2.2. Step 2: Establish Similarity/Dissimilarity Between Networks

This section covers some of the metrics we used to gauge distances between individual networks given the insight they can provide into brain network organizational differences.

#### 2.2.1. KS Statistic

Degree distributions, which help quantify the topology of networks, are likely more similar within distinctive groups than they are between these groups. We employed the log of the KS statistic to quantify this potential dissimilarity.

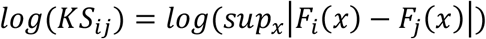

*KS*_*ij*_, a scalar, is the Kolmogorov-Smirnov statistic between nodal degree distributions for individuals *i* and *j. F*_*i*_(*x*) represents the empirical distribution function for observations from the nodal degree vector ***d***_***i***_. So, *sup*_*x*_|*F*_*i*_(*x*) − *F*_*j*_(*x*)| gives the biggest difference between the empirical degree distributions between individuals *i* and *j*. Bigger values indicate more dissimilarity.

*A note on the logarithmic transformation of the KS statistic*: when all distances are nonnegative, it is common practice to take a log transformation. Within our simulations, KS was the only metric which saw improvements in power or type I error when taking such a transformation. For ease of interpretability, none of the other distances presented here utilized a logarithmic transformation.

#### 2.2.2. Jaccard Index

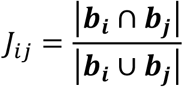

*J*_*ij*_, a scalar, is the Jaccard index between key node vector *i* and key node vector *j*. Since ***b***_***i***_ and ***b***_***j***_ represent the (binary) key node vectors of individuals *i* and *j*, |***b***_***i***_ ∩ ***b***_***j***_| gives the number (count) of nodes that share the same key status between the two networks and |***b***_***i***_ ∪ ***b***_***j***_| gives the total number of nodes. That is, *J*_*ij*_ gives the proportion of nodes that share key status between individuals *i* and *j*. Values of *J*_*ij*_ range from 0 (no overlap) to 1 (perfect overlap--networks are the same).

#### 2.2.3. Euclidean Distance

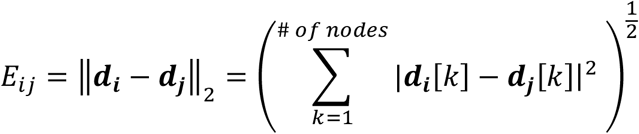

*E*_*ij*_, a scalar, is the Euclidean distance between degree distribution *i* and degree distribution *j*, where ***d***_***i***_[*k*] represents the degree of node *k* for individual *i*. Bigger values of Euclidean distance indicate more dissimilarity.

#### 2.2.4. Minkowski Distance

See Supplemental

#### 2.2.5. Canberra Distance

See Supplemental

### 2.3. Step 3: Evaluating Differences between Networks

#### 2.3.1. Standard F-test

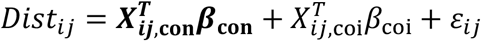

*Dist*_*ij*_ represents the distance between nodal vectors of individuals *i* and *j. Dist*_*ij*_ is a generic placeholder for any metric outlined previously in Step 2, i.e., Jaccard Index (*J*_*ij*_), KS Statistic (*KS*_*ij*_), Minkowski Distance 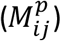, or Canberra Distance (*C*_*ij*_).

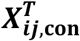, an 1 × (p − 1) vector, contains the intercept and differences in confounding covariates (e.g., for our data, sex, educational attainment, age, and body mass index) between individuals *i* and *j* (with corresponding unknown (*p* − 1) × 1 parameter vector ***β***_**con**_) to control for differences that may confound the relationship between the covariate of interest and the given distance.

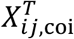, a scalar, contains the difference in the covariate of interest (or an indicator of different group membership for group-based analyses) between individuals *i* and *j* (with corresponding unknown parameter *β*_coi_).

Splitting the design matrix ***X***, an *n* × p matrix, into confounding and of interest covariates is purely a notational preference. 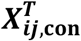 and 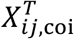 can be combined into the 1 × p vector 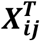 (with corresponding unknown *p* × 1 parameter vector ***β***).

*ε*_*ij*_ accounts for the random error in the distance (Jaccard, KS, etc.) value. If *ε*_*ij*_ were independent, homoscedastic, and approximately normally distributed, the F-test of a standard linear regression would be an appropriate test. We expect correlation among observations from the same individual, so we included this standard testing procedure here mainly for comparison.

As an example, to test whether there is an association between IQ (continuous) and the spatial consistency of hub nodes (top 20% highest degree) after controlling for age (continuous), sex (binary), and treatment (binary) status, our model would be

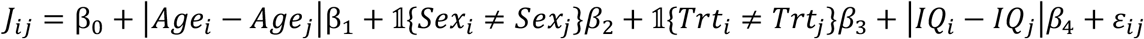

with the associated null hypothesis *H*_0_: β_4_ = 0.

#### 2.3.2. Standard F-test with Individual Level Fixed Effects (F-test with ILE)

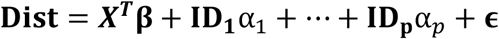

**Dist** is an *n* × 1 vector of known distance metrics (as outlined in Step 2), ***X***^***T***^ is the *n* × p design matrix (intercept optional) of known covariates with corresponding *p* × 1 unknown parameter vector **β, ID**_**i**_ is the *n* × 1 known indicator variable for individual *i* with corresponding unknown parameter α_*i*_. By accounting for individual level effects, we hope to induce independence, homoscedasticity, and normality for the error terms within the *n* × 1 random vector **ϵ**. This would (potentially) allow for an F-test to appropriately evaluate the covariates of interest.

#### 2.3.3. Mixed Model Approach

It should be mentioned that we attempted a mixed model formulation where each individual had their own random effect (where **Dist** = ***X***^***T***^**β** + **ID**_**1**_b_1_ + ⋯ + **ID**_**p**_b_*p*_ + **ϵ** and b_*i*_ *∼ N*(0, *g*) for all *i*). We also attempted b_*i*_ *∼ N*(0, *g*_*i*_) for all *i*. Unfortunately, these calculations were too computationally intensive in the simulations we ran (using the `lmer` function of the `R` package `lme4` (Bates et al., 2015)). Instead, we tried a generalized least squares approach outlined in the next section.

#### 2.3.4. Feasible Generalized Least Squares

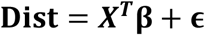

**Dist** and ***X***^***T***^**β** (intercept included) are as before, but instead we assume **ϵ** *∼* (**0, Σ**), where **Σ** is the *n* × n covariance matrix. Generalized least squares (GLS) allows for estimating parameters when there is correlation among the residuals in ordinary least squares regression. However, GLS requires **Σ** to be known. An unrestricted *n* × n covariance matrix has *n*(*n* + 1)/2 parameters to estimate. This is infeasible as we only have *n* observations. Thus, we restricted the form of **Σ** in order to estimate it. For a detailed explanation, please see Supplemental Materials.

#### 2.3.5. Permutation Test

A permutation test requires no knowledge of how the test statistic of interest is distributed under the null hypothesis (e.g., *H*_0_: no significant difference among IQ). The distribution under the null hypothesis is empirically “generated” by permuting data labels. We employed the Freedman-Lane approach (Freedman and Lane, 1983; Kherad-Pajouh and Renaud, 2010) using the `lmperm` function in the `R` package `permuco` (Frossard and Renaud, 2019a) with 10,000 permutations, while permuting across individuals to preserve exchangeability. For a detailed explanation, please see Supplemental Materials as well as the package vignette for `permuco` (Frossard and Renaud, 2019b).

#### 2.3.6. MDMR

Multivariate distance matrix regression (MDMR) is an existing method which has been included here for comparison. It tests the significance of associations of response profile (dis)similarities and a set of predictors. Originally this was done using only permutation (Anderson, 2001), but has been extended to analytic p-values as well (McArtor et al., 2017). MDMR was run using the `mdmr` function in the `MDMR` package in `R` (Mcartor et al., 2018). Both the permutation and analytic versions were run with the *n*_*p*_ × *n*_*p*_ distance matrix ***D*** (the distance matrix analog of **Dist**) and the *n*_*p*_ × *p* design matrix ***X***_***p***_ (covariates of interest for each participant). The permutation method was run with 10,000 permutations.

## 3. Simulation Studies

### 3.1. Data

We conducted four simulation studies to assess how well our proposed approaches could detect various relationships between brain network properties and covariates of interest. Each simulation contained 100 subjects, with four covariates of interest. A fair coin was flipped for each subject to determine their sex (*SEX* = male or female). Half of subjects were assigned to treatment and the other half were assigned to placebo (TRT = treatment or placebo). IQ and Age were both simulated from a normal distribution with mean of 100 and a standard deviation of 15 (rounded to the nearest integer). This resulted in 2 binary (*SEX, TRT*), and 2 continuous (*AGE, IQ*) covariates--variables were given names purely for purposes of explication.

For simulations 1-3, we simulated fMRI connectivity matrices with 268 nodes each to mimic the experimental data detailed in the next section. In each simulation, a 268×268 symmetric matrix (with entries ranging from 0 to 1) was generated for each subject. Entries within each connectivity matrix were drawn from three types of distributions: a) a noise distribution, *Beta*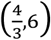; b) a “same-for-all” signal distribution, *Beta*(4,6); and c) a signal distribution dependent on covariates and signal percentage, *Beta*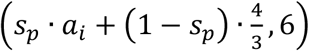 where 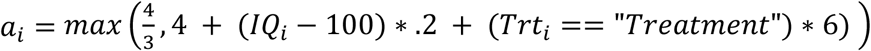 represented the covariate-dependent parameter and *s*_*p*_ represented the signal percentage (from 0 to 100%). When the signal percent (*s*_*p*_) was 100%, 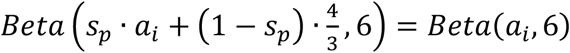. Similarly, when signal percent was 0%, 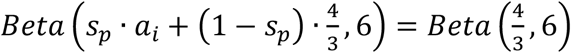, and was therefore identical to the noise distribution and no longer dependent on covariates.

Simulation 1 had a 15×15 region where all individuals had the same signal distribution, a 15×15 region where the signal distribution was dependent on covariates, and the rest of the 268×268 connection matrix was drawn from the noise distribution. Simulation 2 was the same as 1, but the signal distribution dependent on covariates was moved to expand the border of the 15×15 same signal distribution to a combined 21×21 signal region. Simulation 3 was the same as 1, with two additional 15×15 regions where all individuals had the same signal distribution. For a drawn to scale representation of these simulations, see Figure 3. It should be noted that we assumed the connectivity matrices to be symmetric, with 0 entries along the diagonal.

**Figure 3:**
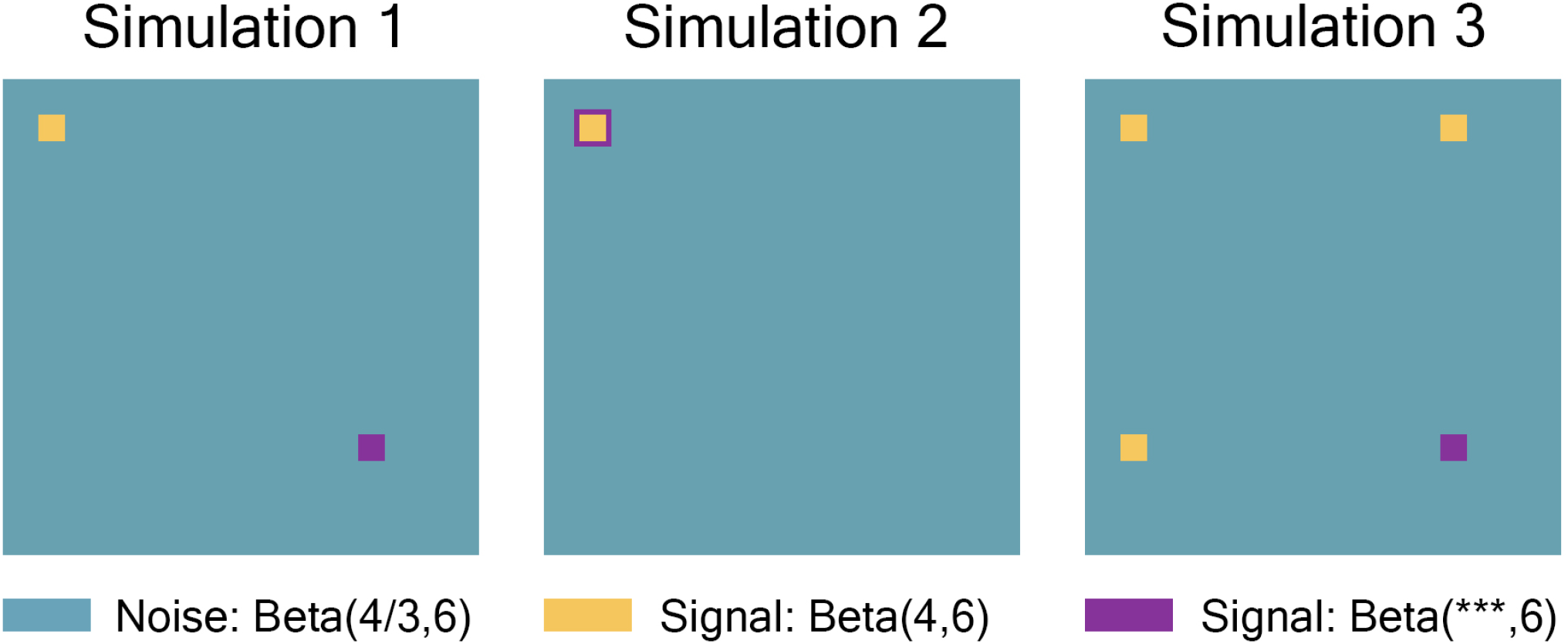
Simulation 1 had a 15×15 region where all individuals had the same signal distribution, a 15×15 region where each the signal distribution was dependent on covariates, and the rest of the 268×268 connection matrix was drawn from the noise distribution. Simulation 2 was the same as 1, but the signal distribution dependent on covariates was moved to expand the border of the 15×15 same signal distribution to a combined 21×21 signal region. Simulation 3 was the same as 1, with two additional 15×15 regions where all individuals had the same signal distribution. The plots were drawn to scale.

Represented by the same colors from Figure 3 simulated connectivity matrices, Figure 4 displays the distributions used for those matrices as signal percentage increased. The noise distribution was distributed Beta(4/3,6) and was shown in teal (not affected by signal percentage). The “same-for-all” signal region was distributed Beta(4,6) and is in yellow (not affected by signal percentage). The “covariate-dependent” signal region can be seen in purple and was distributed Beta(***,6). The *** parameter had some distribution based upon the underlying covariate distribution and signal percentage. There are five purple distributions in each plot, representing the 0.01, 0.25, 0.5, 0.75, and 0.99 quantiles from the *** distribution (for example, the 0.25 quantile distribution is represented by an individual with an IQ of 69 and a “Treatment” status or an individual with an IQ of 99 and a “Placebo” status; the 0.75 quantile distribution is represented by an individual with an IQ of 101 and a “Treatment” status or an individual with an IQ of 131 and a “Placebo” status). Further, the “covariate-dependent” (purple) signal region’s distribution goes from being the same as the noise region’s distribution (at 0% signal) to more and more different than the noise region’s distribution as signal percentage increases. Note that the 0.01 quantile curve overlays the noise curve for all signal intensities.

**Figure 4:**
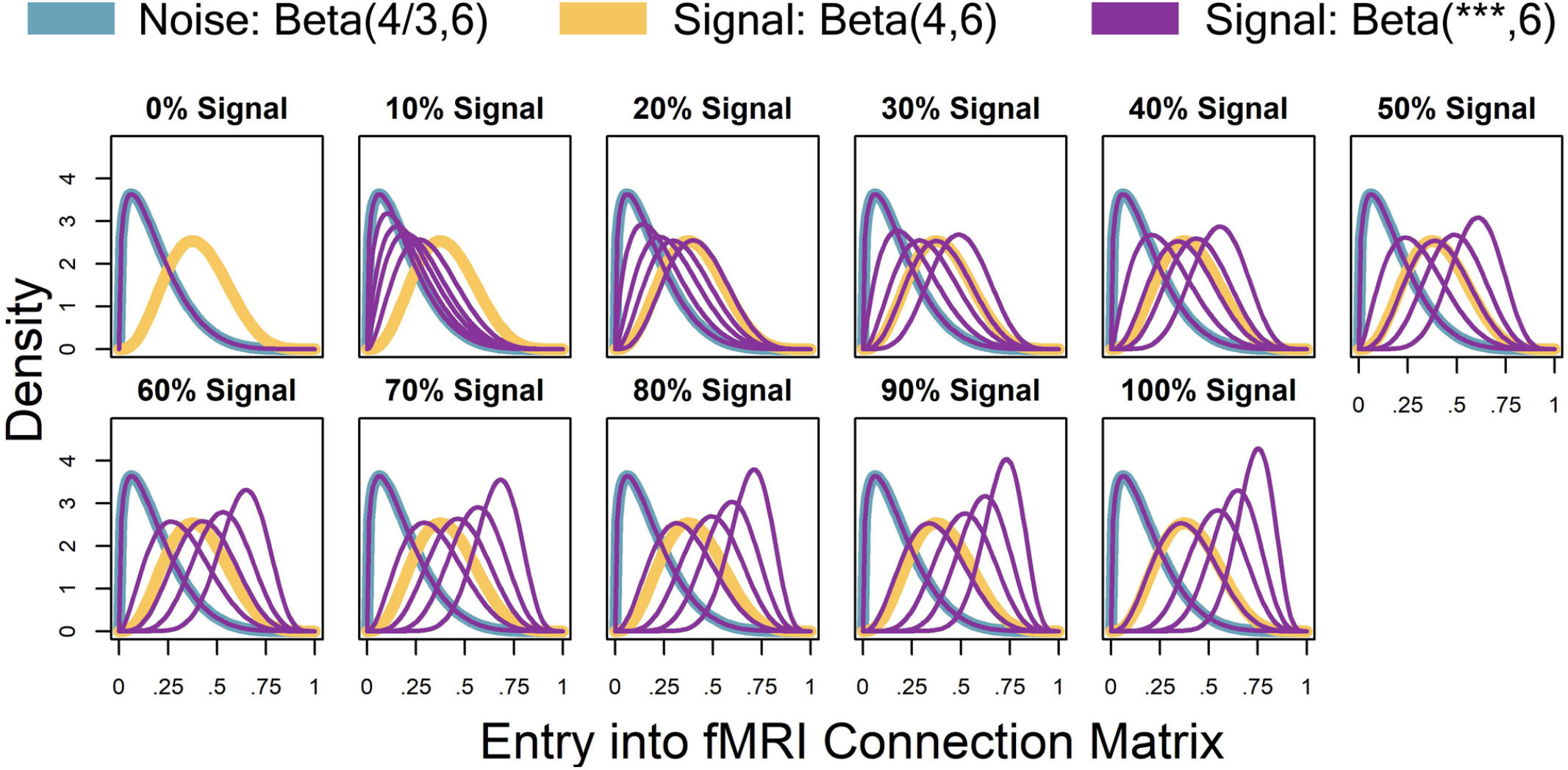
Represented by the same colors from Figure 3 simulated connectivity matrices, Figure 4 displays the distributions used for those matrices as signal percentage increased. The noise distribution was distributed Beta(4/3,6) and was shown in teal (not affected by signal percentage). The “same-for-all” signal region was distributed Beta(4,6) and is in yellow (not affected by signal percentage). The “covariate-dependent” signal region can be seen in purple and was distributed Beta(***,6). The *** parameter had some distribution based upon the underlying covariate distribution and signal percentage. There are five purple distributions in each plot, representing the 0.01, 0.25, 0.5, 0.75, and 0.99 quantiles from the *** distribution (for example, the 0.25 quantile distribution is represented by an individual with an IQ of 69 and a “Treatment” status or an individual with an IQ of 99 and a “Placebo” status; the 0.75 quantile distribution is represented by an individual with an IQ of 101 and a “Treatment” status or an individual with an IQ of 131 and a “Placebo” status). Further, the “covariate-dependent” (purple) signal region’s distribution goes from being the same as the noise region’s distribution (at 0% signal) to more and more different than the noise region’s distribution as signal percentage increases. Note that the 0.01 quantile curve overlays the noise curve for all signal intensities.

For simulation 4, we simulated nodal degree vectors of length 268 (instead of 268 x268 connectivity matrices as in simulations 1-3) to assess the method’s ability to detect distributional differences rather than location differences. All entries of each individual’s degree vector were simulated Normal(100 + *s*_*p*_ · *a*_*i*_, 1), where *s*_*p*_ (signal percent) and *a*_*i*_ (covariate-dependent parameter for individual *i*) were as described with simulations 1-3. Within Figure 5, there were five purple distributions in each plot, representing the 0.01, 0.25, 0.5, 0.75, and 0.99 quantiles from the distribution of the mean parameter, 100 + *s*_*p*_ · *a*_*i*_. At 0% signal, all individual’s nodal degree vectors were drawn from *Normal*(100,1). As signal percentage increased, the mean parameter of the covariate-dependent distributions became more and more distinct.

**Figure 5:**
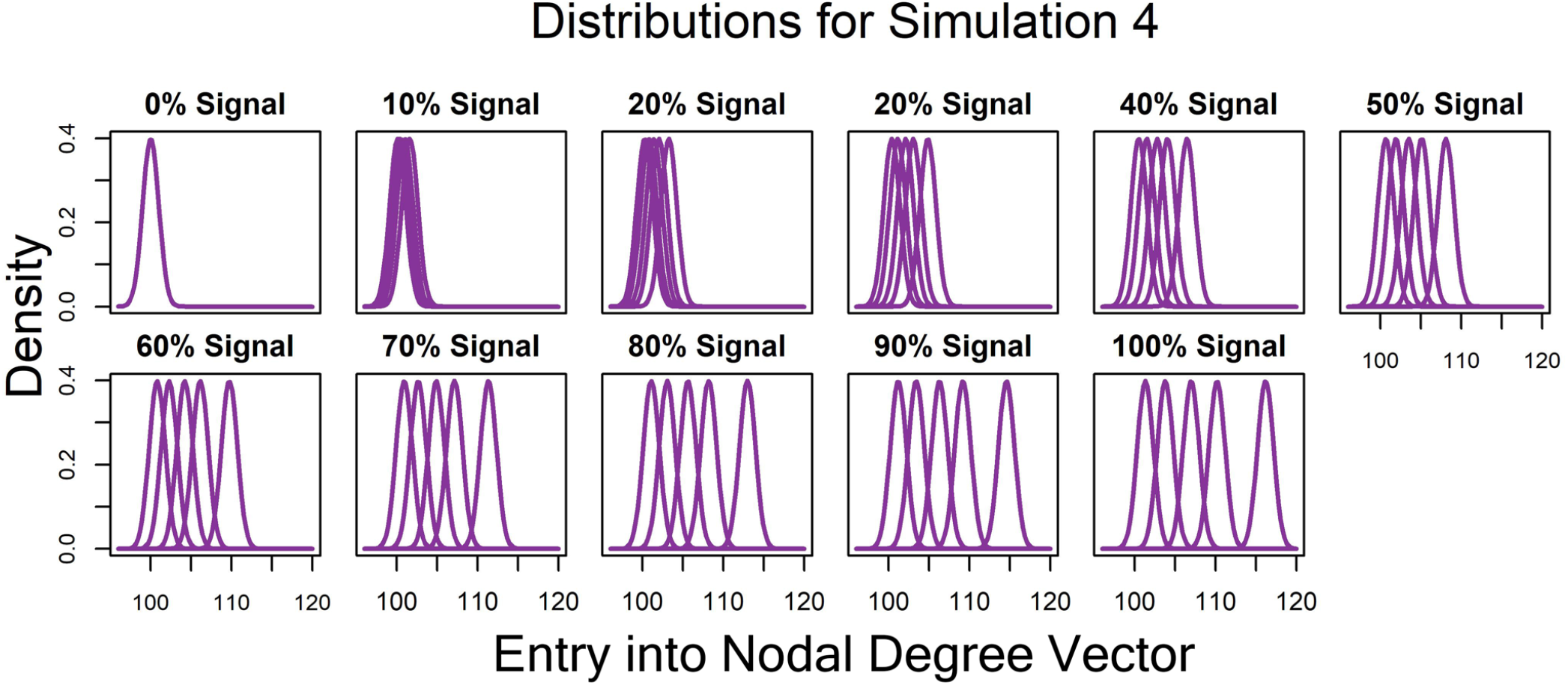
For simulation 4, we simulated nodal degree vectors of length 268 (instead of 268 x268 connectivity matrices as in simulations 1-3) to assess the method’s ability to detect distributional differences rather than location differences. All entries of each individual’s degree vector were simulated Normal (100 + *s*_*p*_ · *a*_*i*_, 1), where *s*_*p*_ (signal percent) and *a*_*i*_ (covariate-dependent parameter for individual i) were as described with simulations 1-3. There were five purple distributions in each plot, representing the 0.01, 0.25, 0.5, 0.75, and 0.99 quantiles from the distribution of the mean parameter, 100 + *s*_*p*_ · *a*_*i*_. At 0% signal, all individual’s nodal degree vectors were drawn from Normal (100,1). As signal percentage increased, the mean parameter of the covariate-dependent distributions became more and more distinct.

### 3.2. Results

We assessed methods with the four simulation scenarios detailed in the previous section. Each simulation was run 10,000 times.

Nodal degree vectors (used for the KS and Euclidean distances) were created by summing across rows of the connectivity matrices. Key nodes of interest (binary degree vectors used for the Jaccard index) based on node degree were identified, selecting the top 20% highest degree (hub) nodes and mapping those to 1 while mapping all remaining nodes to 0. The KS statistic and Euclidean distance were calculated for each pair of individuals using their nodal degree vectors. The Jaccard distance was calculated for each pair of individuals using their binary degree vectors.

The percentages of p-values less than *α* = 0.05 for the covariates of interest were recorded for each combination of signal percent (0%, 10%, …, 100%), distance metric (KS, Jaccard, Euclidean), and testing framework (F-test, Permutation, GLS, F-test with individual level effects, MDMR Analytic and MDMR Permutation). In this section, we discuss whether type I error rate was controlled and at what signal percent the 80% power threshold was reached. Then, we compare results with MDMR. For a visual display of the results, see Figure 6. For more plots and a visual display with additional distance metrics, please see Supplemental Materials.

**Figure 6:**
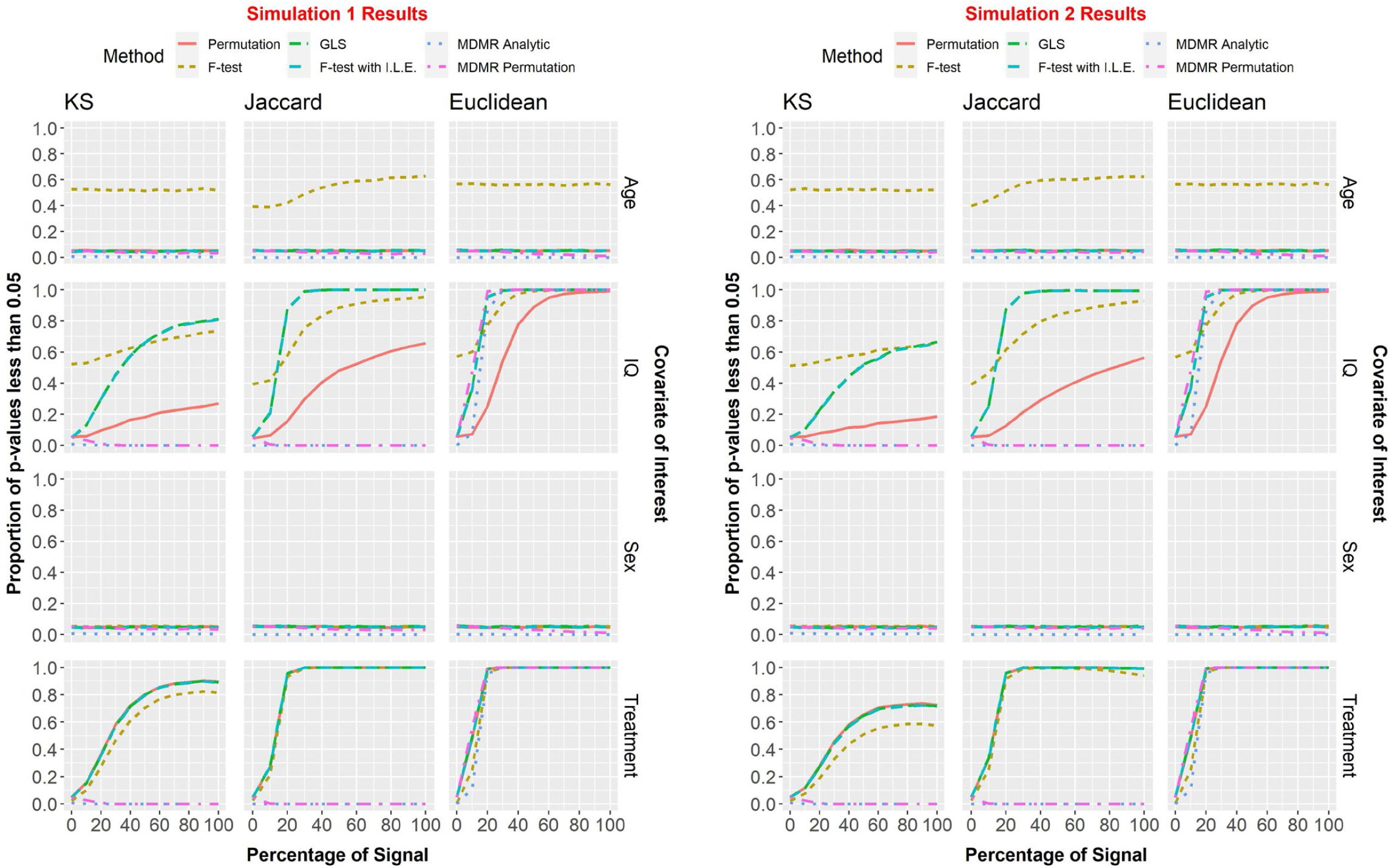
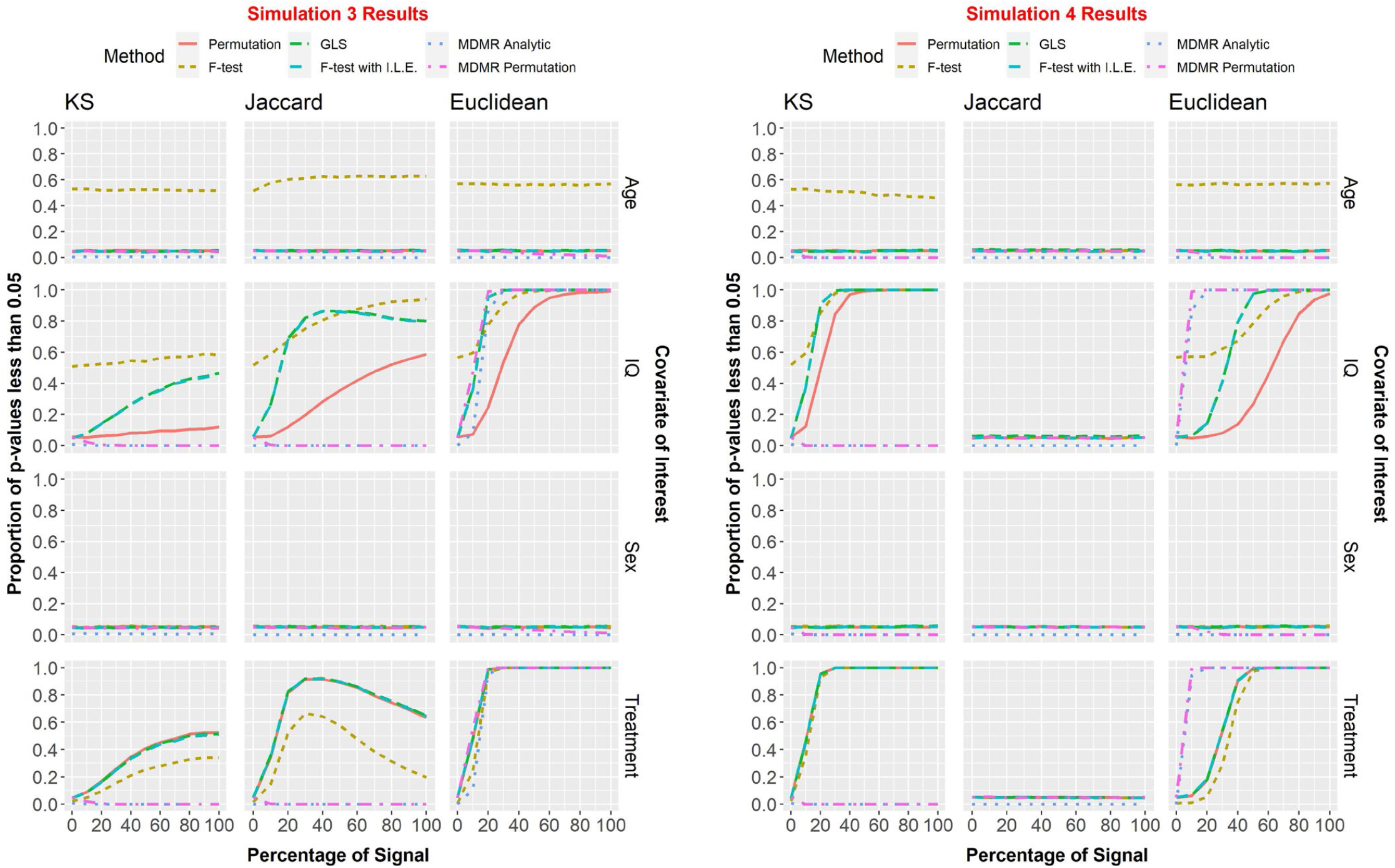
We assessed methods with the four simulation scenarios detailed in the previous section. Each simulation was run 10,000 times. The percentages of p-values less than *α = 0.05* for the covariates of interest were recorded for each combination of signal percent (0%, 10%, …, 100%), distance metric (KS, Jaccard, Euclidean), and testing framework (F-test, Permutation, GLS, F-test with individual level effects, MDMR Analytic and MDMR Permutation).

The standard F-test did not control type I error when testing Age. It was included in the Figure for reference, but is not mentioned any further in this section.

#### 3.2.1. KS

All methods (not including F-test) adequately controlled type I error. <u>Simulation 1:</u> The GLS and F-test with ILE methods reached 80% power within 100% of signal for continuous covariates. The Permutation method never reached 80% power for continuous covariates. For binary covariates, all four methods reached 80% power within 60% of signal. <u>Simulations 2-3</u>: No methods reached 80% power at any level of covariate-dependent signal. <u>Simulation 4</u>: The GLS and F-test with ILE methods reached the power threshold within 20% of signal. Permutation reached the threshold within 30% of signal. All three methods controlled type I error.

#### 3.2.2. Jaccard

All methods (not including F-test) adequately controlled type I error. <u>Simulations 1-3, IQ</u>: The GLS and F-test with ILE methods reached the power threshold within 20-30% of signal; the permutation method never reached the minimum power requirement. <u>Simulations 1-3, Treatment</u>: The GLS, F-test with ILE, and Permutation methods reached the power threshold within 20% of signal. **Surprisingly, as signal percentage increased, some situations had decreasing power with some even falling far back below the power threshold. This only occurred with the Jaccard distance metric**. <u>Simulation 4</u>: No methods could detect significance. All methods’ power levels hovered around *α* = 0.05, the significance level.

#### 3.2.3. Euclidean

All methods (not including F-test) adequately controlled type I error. <u>Simulations 1-3, IQ</u>: The GLS and F-test with ILE methods reached the power threshold within 20% signal. The permutation method reached the minimum power requirement within 50% of signal. <u>Simulations 1-3, Treatment</u>: All methods reached the power threshold within 20% signal. <u>Simulation 4, IQ</u>: The GLS and F-test with ILE methods reached the power threshold within 50% signal. The permutation method reached the minimum power requirement within 80% of signal. <u>Simulation 4, Treatment</u>: All methods reached the power threshold within 40% signal.

#### 3.2.4. Comparison with MDMR

<u>KS and Jaccard</u>: Neither MDMR Permutation nor MDMR analytic had adequate power for the KS and Jaccard distances. Our approaches vastly outperformed them for all simulation scenarios except for Simulation 4/Jaccard in which the results were comparable. <u>Euclidean</u>: For Simulations 1-3, the MDMR permutation method had comparable power with the GLS and F-test with ILE methods, while MDMR analytic had slightly less power. For Simulation 4, both MDMR methods had considerably more power than any of the methods within our framework. MDMR analytic Type I error rate remained close to 0 for all simulations. MDMR permutation started off with the expected Type I error rate of approximately 0.05, but as signal increased, Type I error rate approached 0 as well.

## 4. Experimental Studies

### 4.1. Data

The HCP data released to date include 1,200 individuals (Van Essen et al., 2013). Of those, 1,113 (606 females; 283 minority) have complete MRI images, cognitive testing, and detailed demographic information. The 397 subjects used are what remained after quality control assessment of head motion and global signal changes for both scan types, removal of those with missing data, and random selection of one individual from each family to ensure between-subject independence. Participants in the HCP completed two resting-state scans and two working memory scans. The two scans were collected with different phase encoding (right to left vs. left to right). The resting-state scans were collected back-to-back while participants quietly viewed a fixation point. The 2-back task was a block design that interleaved the 2-back condition with a 0-back condition and a rest period. The working memory task utilized photos and different blocks had different photo types (faces, body parts, houses, tools). Participants were alerted prior to each block to indicate the task type. For the 2-back they were instructed to respond anytime the current stimulus being presented matched the stimulus two trials back. The HCP performed extensive testing and development to ensure comparable imaging at the two sites (Van Essen et al., 2012). The blood oxygenation level dependent (BOLD)-weighted images were collected using the following parameters: TR = 720 ms, TE = 33.1 ms, voxel size 2 mm^3^, 72 slices, 1,200 images.

The main covariate of interest for this analysis was Fluid Intelligence. Other covariates included in model formulation were Age, BMI, Education, Handedness, Income, Alcohol Abuse, Alcohol Dependence, Ethnicity, Race, Sex, and Smoking Status. For a summarization and explanation of the variables, see Tables 1 and 2.

**Table 1:** Summarization and explanation of HCP covariates treated as continuous (within the regression framework).

|  | Mean<br>(S.D.) | Notes |
| --- | --- | --- |
| Age | 28.7 (3.7) | In Years. |
| BMI | 26.5 (5.1) | Body Mass Index. |
| Education | 15.0 (1.7) | Integer Values 11 to 17 (years pf education completed). |
| Fluid Intelligence | 17.3 (4.7) | Integer valued from 4 to 24. |
| Handedness | 65.1 (43.6) | Values range from -100 to 100 by 5 (-100, -95, ..., 95, 100) |
| Income | 5.0 (2.1) | SSAGA income score - Total household income:<br><\$10,000 = 1, 10K-19,999 = 2, 20K-29,999 = 3, 30K-39,999 = 4, 40K-49,999 = 5, 50K-74,999 = 6, 75K-99,999 = 7, >=100,000 = 8. |

**Table 2:**
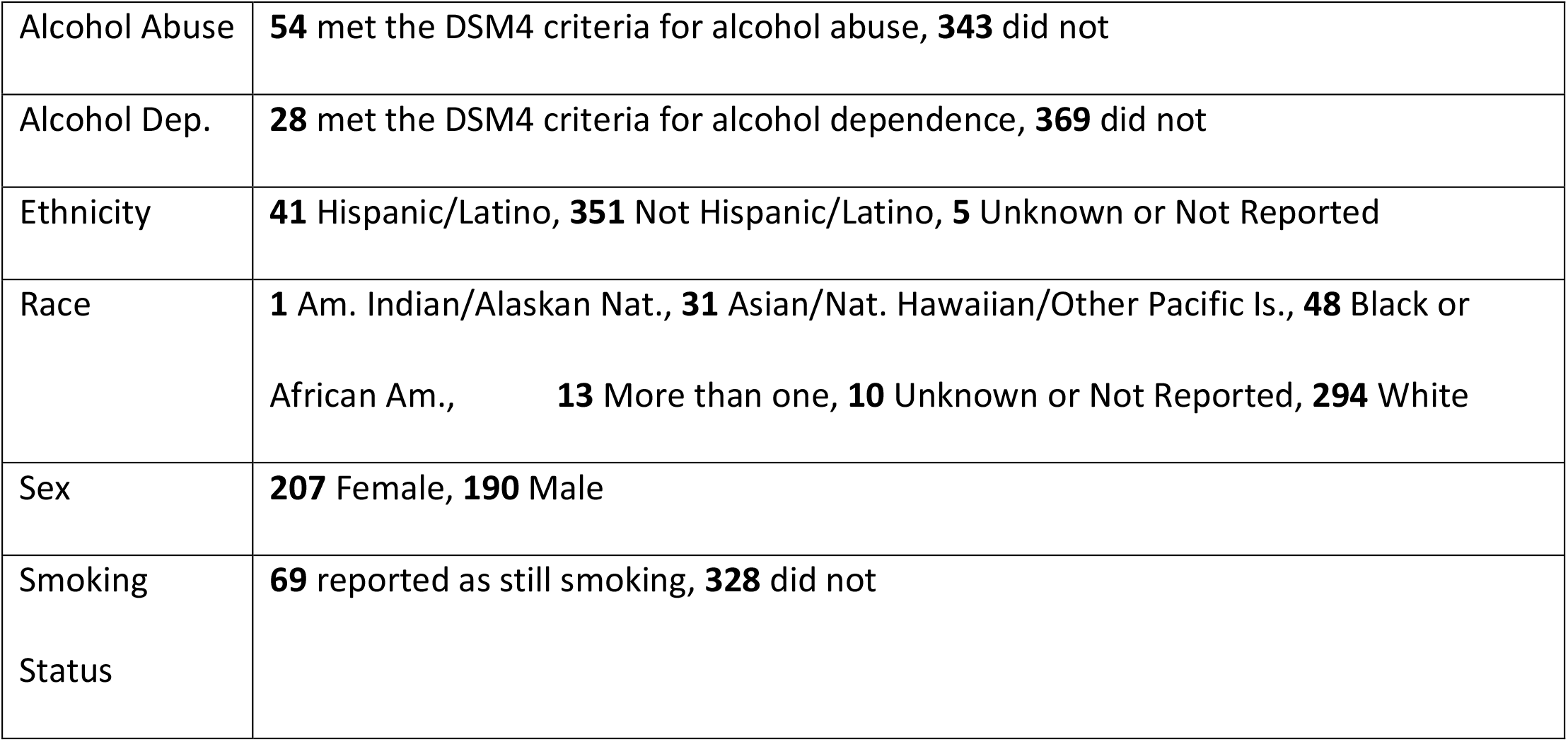
Summarization and explanation of HCP covariates treated as categorical (within the regression framework).

### 4.2. Data Processing and Network Generation

The current project used the minimally processed fMRI data provided by the HCP (Glasser et al., 2013) for resting state and working memory. Additional preprocessing steps included: motion correction using ICA-AROMA (Pruim et al., 2015), removal of the first 14 volumes from each scan, and band-pass filtering (0.009– 0.08 Hz) to remove physiological noise and low-frequency drift. It was necessary to account for the block design of the working memory task. First, the block design was modeled in SPM12, yielding regressors for the 0-back and rest blocks as well as the cues. Additionally, since each scan was collected twice with different phase encoding, the scans were concatenated and a scan-specific regressor was added. All regressors, including the total gray matter (GM), white matter (WM), and cerebrospinal fluid (CSF) signals and realignment parameters were used in a single regression analysis. The residual signal following regression of the extraneous variables was retained only for periods aligned with the 2-back blocks. The blocks of data were then concatenated into a single time series containing 274 BOLD images. Resting state data followed a similar pipeline, but without the task-specific regressors. The resulting resting state data contained 2,372 bold images. After preprocessing, the brain was parcellated into 268 regions as defined in the Shen Atlas (Shen et al., 2013) and the signal from all voxels within each region was averaged for each participant. A functional network was constructed for each participant by computing the Pearson (full) correlation between the resultant time series for each region pair. Negative correlation values were set to zero because the neurobiological interpretation of positive and negative edges are very different (Parente et al., 2017; Schwarz and McGonigle, 2011). In addition, since the distributions of network variables (such as degree) are different for positive and negative edges (Fraiman et al., 2009; Saberi et al., 2021). Although negative correlations are not regularly used in network neuroscience, if they are used, positive and negative networks should be generated and assessed separately (Schwarz and McGonigle, 2011). For the current work we focused on positive networks, but one could simply perform an additional analysis on negative networks if so desired.

### 4.3. Results

Nodal degree vectors (used for the KS and Euclidean distances) were created by summing across rows of the connectivity matrices. Key nodes of interest (binary degree vectors used for the Jaccard index) based on node degree were identified, selecting the top 20% highest degree (hub) nodes and mapping those to 1 while mapping all remaining nodes to 0. KS statistic and Euclidean distance were calculated for each pair of individuals using their nodal degree vectors. The Jaccard distance was calculated for each pair of individuals using their binary degree vectors.

Distance covariates for each pair of individuals were calculated. A continuous variable’s distance (Age, for instance) was calculated as |*Age*_*i*_ − *Age*_*j*_| for the pair of individuals *i* and *j*. A binary or categorical variable’s distance (Education, for instance) was calculated as 𝟙{*Edu*_*i*_ ≠ *Edu*_*j*_} for the pair of individuals *i* and *j*.

We evaluated differences between networks using the Standard F-test with ILE, as it was the best in our simulations at controlling type I error and providing sufficient power across all distance metrics (while also being computationally inexpensive). Parameter and standard error estimates can be found in Supplemental Materials in Tables S1 and S2. Each parameter estimate represented the average amount the given brain distance metric (KS, Jaccard, etc.) changed based on a one-unit difference in the respective covariate, after controlling for other covariates. A complete list of p-values for both resting state and working memory can be seen in Table 3. There have been no adjustments for multiple comparisons.

**Table 3:** P-values for HCP resting state and working memory brain scans when modeled with our given regression framework and tested using the standard F-test with fixed individual level effects. Parameter estimates and standard errors can be found in Supplementary Materials.

|  | Resting State |  |  | Working Memory |  |  |
| --- | --- | --- | --- | --- | --- | --- |
|  | KS | Jaccard | EUC | KS | Jaccard | EUC |
| Fluid Intelligence | 8.66E-09 | 5.96E-03 | 1.12E-09 | 2.43E-01 | 3.47E-02 | 2.80E-02 |
| Age | 4.52E-03 | 6.35E-02 | 9.09E-05 | 8.71E-02 | 9.68E-04 | 9.41E-09 |
| Alcohol Abuse | 8.55E-05 | 4.35E-01 | 1.10E-02 | 7.78E-01 | 2.02E-01 | 4.45E-01 |
| Alc. Dependence | 3.03E-01 | 2.26E-01 | 2.73E-01 | 7.85E-01 | 7.84E-02 | 1.86E-01 |
| BMI | 1.26E-01 | 4.97E-03 | 1.40E-05 | 1.43E-02 | 1.62E-01 | 3.95E-01 |
| Education | 1.48E-01 | 3.10E-01 | 4.06E-01 | 9.11E-03 | 2.93E-04 | 2.03E-02 |
| Ethnicity | 8.81E-01 | 4.25E-01 | 4.66E-01 | 7.16E-01 | 1.44E-01 | 4.44E-01 |
| Gender | 4.91E-02 | 6.24E-16 | 4.08E-16 | 7.24E-01 | 8.70E-10 | 1.03E-08 |
| Handedness | 5.17E-01 | 2.43E-02 | 1.13E-01 | 3.70E-01 | 2.30E-01 | 7.14E-01 |
| Income | 3.80E-01 | 2.06E-02 | 1.82E-01 | 7.08E-01 | 6.00E-01 | 3.70E-01 |
| Race | 5.03E-03 | 2.14E-04 | 3.79E-13 | 3.08E-01 | 1.34E-07 | 6.05E-03 |
| Smoking Status | 2.23E-01 | 7.16E-01 | 3.00E-01 | 5.75E-01 | 6.37E-01 | 8.63E-01 |

After adjusting for the other confounding variables, the covariate of interest, Fluid Intelligence, had a statistically significant relationship with all distance metrics (KS, Jaccard, Euclidean) for Resting State fMRI, and non-KS distance metrics for Working Memory fMRI. The level of significance was considerably higher for the resting-state data. Figure 7a shows that the resting-state spatial distribution of brain hubs are relatively concentrated in the brain regions making up the default mode network (known to have high degree at rest). However, there is a higher concentration of hubs localized to the default mode network in the bottom quartile of fluid intelligence compared to those in the top quartile. Figure 7b shows that hub locations during the working memory task shift from default mode areas to regions that make up the central executive attention network (CEN) more so in the top quartile of fluid intelligence. Note that in those individuals in the bottom quartile of intelligence, the hubs are most prominent in the areas of the default mode network even during the working memory task, though the magnitude of the overlap is lower. The CEN maps onto what is sometimes referred to as the frontoparietal network. There is evidence that the density of structural connectivity in that network predicts WM capacity, with higher capacity individuals exhibiting greater connectivity (Ekman et al., 2016).

**Figure 7:**
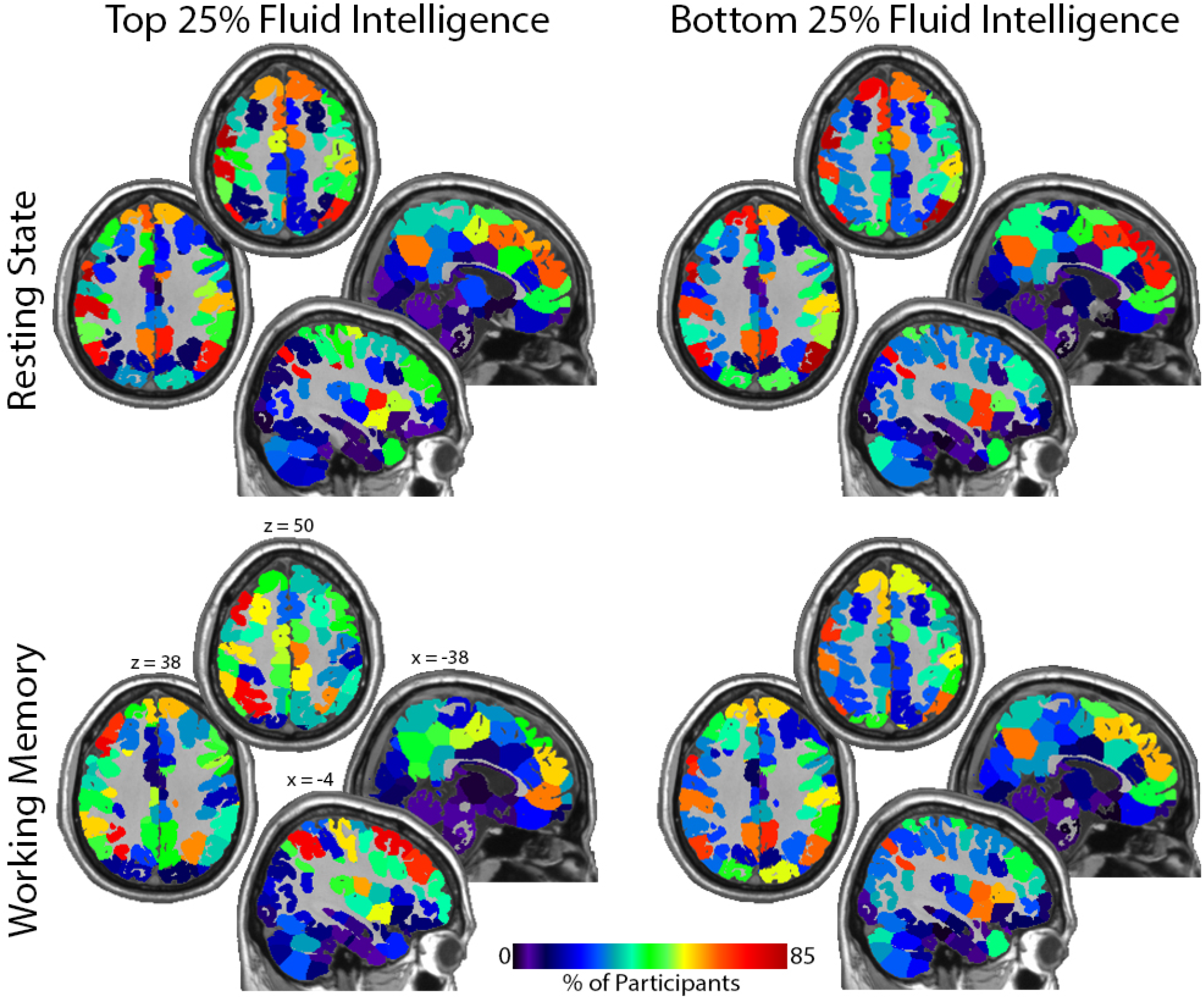
Maps showing the location of network hubs (top 20% degree) for the top and bottom intelligence quartiles. Resting State: note the high incidence of hubs in the medial and lateral parietal cortex and the medial frontal lobe in both groups. These regions are central components of the default mode network. The significant association between degree location and intelligence is demonstrated by the reduction in default mode network hubs with increases in fluid intelligence. Working Memory: Hubs are more concentrated in the central executive network as intelligence increases. This shift is best appreciated in the top quartile group where hubs are concentrated in lateral rather than medial frontal regions and in more superior parietal areas. Note that the hubs remain fairly localized to the default mode network for the bottom quartile group. Each quadrant shows 2 axial and 2 sagittal images. The Montreal neurological Institute (MNI) coordinates shown in the bottom left quadrant apply to all quadrants. Calibration bar applies to all images.

Among the confounding variables, some had significant relationships with all distance metrics except for the Jaccard Index (Age and Alcohol Abuse within Resting State). Ideally, we would expect Jaccard to be statistically significant if Euclidean norms were heavily significant. This could have been due to the Jaccard-related issues that showed up in our simulations. There were also certain variables that were significant for one fMRI task but not the other (Alcohol Abuse, BMI and Education). Some covariates were significant for all location-specific metrics but not for the distance between distribution comparator KS statistic (BMI – Rest; Gender – Rest and WM; Age, Fluid Intelligence and Race – WM). This was indicative of a detectable difference in location-specific degree, but not in distribution.

In addition to our primary analysis detailed above, we also examined these relationships with respect to differences in modularity between individuals employing Scaled Inclusivity given that modularity analyses can capture the spatial distribution of intrinsic brain networks that are associated with various cognitive tasks (Moussa et al., 2012). The description of this secondary analysis, along with the corresponding results and figures can be found in Supplementary Material.

## 5. Discussion

It is of great interest to compare differences in fMRI connection matrices between individuals by covariates. Our previous work developed a novel permutation testing framework to detect differences between two groups (Simpson et al., 2013b). Here we advanced that work to allow both assessing differences by continuous phenotypes and controlling for confounding variables. We proposed an innovative regression framework to relate distances between brain network features to functions of absolute differences in continuous covariates and indicators of difference for categorical variables, and explored several similarity metrics for comparing distances between connection matrices. The KS statistic measures how different distributions of topological properties vary between two individuals. Key-node metrics (like the Jaccard Index) quantify how much the spatial location of key brain network regions differs between two networks. The Euclidean norm (along with Canberra, Minkowski, Weighted Jaccard, etc.) measures whether the spatial location of degree-weighted brain network regions differ. Several additional similarity/dissimilarity metrics mentioned below were also assessed, but details were left out for the sake of brevity. The binary network-based metrics left out included the Russel-Rao, Kulczynski, and Baroni-Urbani and Buser. Little difference or improvement was found (in simulations 1-3) for any of these metrics when compared with the commonly used Jaccard Index. The weighted network-based metrics left out included the Weighted Jaccard, Manhattan, and Maximum distances, with none different or better than the already presented Canberra or Minkowski distances (when tested in simulations 1-3). Any other distance metrics could be used. Future work might include: 1) testing other metrics, for example, Distance Correlation (Székely et al., 2007; Székely and Rizzo, 2009); 2) further investigating the behavior of the Jaccard metric; and 3) taking a deeper dive into understanding when and how to choose a distance metric.

Several standard methods for estimation and inference were adapted to fit into our regression framework: Standard F-test, F-test with individual level effects (ILE), Generalized Least Squares (GLS), and Permutation (inference only). All combinations of these approaches and the distance metrics were assessed via four simulation scenarios. The KS statistic was found to have low comparable power when testing location-specific differences (relative to the other distance metrics tested). However, if interested in comparing distributions, the KS statistic was preferred. The Jaccard Index did not have consistent or predictable power, and further work should investigate the reasons underlying this. An argument could be made for several different distance metrics when detecting location-specific differences between degree vectors, as they had very similar results. We prefer to use Euclidean distance, as it is commonly recognized and had the best overall (slightly) type I control and power. Future work could include a further investigation into how and when to choose specific distance metrics. Regarding the comparison of estimation and testing methods (Standard F-test, F-test with ILE, GLS, Permutation), we recommend the F-test with ILE as it was among the best at controlling type I error and providing sufficient power while also being computationally inexpensive.

An FGLS approach for estimation and inference was proposed in this framework. Although it is slower (computationally) than the Standard F-test with ILE, the FGLS approach performed just as well as the recommended Standard F-test with ILE (in terms of type I and type II error).

An analysis of the HCP data was completed using the Standard F-test with ILE and several distance metrics. After adjusting for the other confounding variables, the covariate of interest, Fluid Intelligence, was significantly related with all distance metrics (KS, Jaccard, Euclidean) for both fMRI tasks (Resting State, Working Memory). This analysis suggested the hubs become more strongly concentrated in default mode network regions at rest with decreasing fluid intelligence. As intelligence increases, there are greater shifts in the spatial locations of hubs from the default mode network at rest to the central executive network during a working memory task.

Many existing methods exist for relating network metrics and phenotypes. Such methods include, but are not limited to traditional network models (e.g., exponential random graph models (Lehmann et al., 2021; Simpson et al., 2012, 2011)), tensor regression works on brain network (e.g. (Zhang et al., 2018, 2019)), Bayesian approaches (e.g. (Dai and Guo, 2017; Wang et al., 2017)), and statistical learning techniques (RC et al., 2015; Varoquaux and Craddock, 2013; Xia et al., 2020). We believe our method most closely relates to Multivariate Distance Matrix Regression (MDMR). MDMR tests the significance of associations of response profile (dis)similarities and a set of predictors. Originally this was done using only permutation (Anderson, 2001), but has been extended to analytic p-values as well (McArtor et al., 2017). Via simulation, a comparison to our regression framework showed that MDMR performs relatively well for the Euclidean metric (as well as other Minkowski distances—see Supplemental) in terms of power. However, it’s type I error rate properties were not well understood. For many covariates, close to 0% of tests had p-values less than 0.05. Further investigation should be done to better understand this property. Further, MDMR was not able to detect differences on either the KS or Jaccard Metrics. Our framework vastly outperformed MDMR for these two metrics for all simulation scenarios except for Simulation 4/Jaccard in which the results were comparable.

We have developed a testing framework that detects whether the spatial location of key brain network regions and distributions of topological properties differ by phenotype (continuous and discrete) after controlling for confounding variables in static networks. More generally, this framework allows relating distances between networks (e.g., Jaccard, K-S distance) to covariates of interest. Our chosen method, F-test with ILE, is computationally inexpensive, generally interpretable, and well understood by most scientists. We believe this adds a convenient tool to the neuroscience toolbox. Future work plans to extend this approach by creating a dynamic network analog that uncovers whether within- and across-task time-varying changes in these spatial and distributional patterns differ by phenotype.

## Acknowledgements

The authors thank Hongtu Zhu, Professor of Biostatistics at UNC Chapel Hill, for suggesting that we assess the logarithmic transformations of the distance metrics. We also thank Dale Dagenbach, Professor of Psychology at Wake Forest University, for his insights into interpreting our HCP results regarding IQ.

## Data Availability Statement

Simulation and HCP code is available: https://github.com/applebrownbetty/braindist_regression. HCP data is publicly available for download.

