## Supplemental for "A Regression Framework for Brain Network Distance Metrics"

### Title

### Supplemental

### 2. Methods

#### 2.2. Step 2: Establish Similarity/Dissimilarity Between Networks

##### *2.2.4. Minkowski Distance ($L^p$ norm distance)*

$$M_{ij}^p = \left\|d_i - d_j\right\|_p = \left( \sum_{k=1}^{\#\ of\ nodes} |d_i[k] - d_j[k]|^p \right)^{\frac{1}{p}}$$

$M_{ij}^p$ is the Minkowski distance of order $p$ between degree distribution $i$ and degree distribution $j$, where $d_i[k]$ represents the degree of node $k$ for individual $i$. There are an infinite number of choices of Minkowski distances (the Manhattan distance when $p = 1$, the Euclidean distance when $p = 2$, etc.). Bigger values of Minkowski distances indicate more dissimilarity.

*2.2.5. Canberra Distance*

$$C_{ij} = \sum_{k=1}^{\#\,of\,nodes} \frac{|d_i[k] - d_j[k]|}{|d_i[k]| + |d_j[k]|}$$

$C_{ij}$ is the Canberra distance between degree distribution $i$ and degree distribution $j$, where $d_i[k]$ represents the degree of node $k$ for individual $i$. The Canberra Distance is a weighted version of the Manhattan distance (the Minkowski distance when $p = 1$). Bigger values of Canberra distances indicate more dissimilarity.

#### 2.3. Step 3: Evaluating Differences between Networks

*2.3.5. Feasible Generalized Least Squares (Expanded)*

$$\mathbf{Dist} = \boldsymbol{X^T}\boldsymbol{\beta} + \boldsymbol{\epsilon}$$

$\mathbf{Dist}$ and $\boldsymbol{X^T}\boldsymbol{\beta}$ are as before, but instead we assume $\boldsymbol{\epsilon} \sim (\mathbf{0},\ \boldsymbol{\Sigma}),$ where $\boldsymbol{\Sigma}$ is the $n\ \times \mathrm{n}$ covariance matrix. Generalized least squares (GLS) allows for estimating parameters when there is correlation among the residuals in ordinary least squares regression. However, GLS requires $\boldsymbol{\Sigma}$ to be known. An unrestricted $n\ \times\ \mathrm{n}$ covariance matrix has $n(n+1)/2$ parameters to estimate. This is infeasible as we only have $n$ observations. Thus, we restricted the form of $\boldsymbol{\Sigma}$ in order to estimate it.

Here we propose an artful way to estimate the covariance matrix. Let $\boldsymbol{\Sigma} = \boldsymbol{\sigma^2}\mathbf{I_n} + \boldsymbol{\tau^2}\mathbf{ID_{mat}}\mathbf{ID_{mat}^{\mathit{T}}}$, where $\mathbf{I_n}$ is the $n\ \times \mathrm{n}$ identity matrix and $\mathbf{ID_{mat}} = [\mathbf{ID_1} \mid \mathbf{ID_2} \mid \cdots \mid \mathbf{ID_p}]$ is an $n\ \times p$ matrix such that each $\mathbf{ID_i}$ is the $n\ \times 1$ indicator variable for individual $i$. Now, in order to estimate $\boldsymbol{\Sigma}$, we just need to estimate two parameters: $\sigma^2$ and $\tau^2$. Taking a closer look at $\boldsymbol{\Sigma}$, we notice that each diagonal element is $\sigma^2 + 2\tau^2$, some off-diagonal elements are $\tau^2$, and all other off-diagonal elements are $0$.

Since each diagonal element, $\sigma^2 + 2\tau^2$, is the same, we can estimate it using sample variance:

$$\widehat{\sigma^2 + 2\tau^2} = \frac{1}{n}\left(\mathbf{Dist} - \boldsymbol{X_i^T}\beta\right)^T\left(\mathbf{Dist} - \boldsymbol{X_i^T}\beta\right)$$

Since we have that some off-diagonals of $\boldsymbol{\Sigma}$ are $\tau^2$ and the rest are $0$, each nonzero off-diagonal is the same value. Thus, we can estimate $\tau^2$ using the *idea* of sample covariance:

$$\widehat{\tau^2} = \frac{1}{m}\left(\mathbf{Dist} - \boldsymbol{X}_i^T\beta\right)^T\left(\mathbf{ID_{mat}}\mathbf{ID_{mat}^{\mathit{T}}} - \mathbf{2}\cdot\mathbf{I_n}\right)\left(\mathbf{Dist} - \boldsymbol{X}_i^T\beta\right)$$

where $m$ represents the sum of all entries in $\mathbf{ID_{mat}}\mathbf{ID_{mat}^{\mathit{T}}} - \mathbf{2}\cdot\mathbf{I_n}$, which gives us the number of non-zero off-diagonal entries in the variance-covariance matrix.

Next, we pre-multiply our original model with $\mathbf{L^{-1}}$, the inverse of Cholesky decomposition of $\boldsymbol{\Sigma}$ such that $\boldsymbol{\Sigma} = \mathbf{LL^T}$:

$$\mathbf{L^{-1}Dist} = \mathbf{L^{-1}}\boldsymbol{X^T}\boldsymbol{\beta} + \mathbf{L^{-1}}\boldsymbol{\epsilon} \xrightarrow{yields} \mathbf{Dist}^* = \boldsymbol{X^{*T}}\boldsymbol{\beta} + \boldsymbol{\epsilon}^*$$

where $\boldsymbol{\epsilon}^* = \mathbf{L^{-1}}\boldsymbol{\epsilon} \sim (\mathbf{L^{-1}}\cdot\mathbf{0_n}, \mathbf{L^{-1}\Sigma L^{-T}}) = (\mathbf{0}_n, \boldsymbol{I}_n)$.

We should note here that FGLS calls for the above process to be iterated until convergence (repeat the above steps using $\mathbf{Dist}^*$ and $\boldsymbol{X}^*$). In this case, the method converges adequately in one step, and further iterations are not necessary.

Finally, since post-transformation it is assumed entries of $\boldsymbol{\epsilon}$ are independent, homoscedastic, and approximately normally distributed, we can apply the F-test.

##### *2.3.6. Permutation Test (Expanded)*

A permutation test requires no knowledge of how the test statistic of interest is distributed under the null hypothesis (e.g., $H_0$: no significant difference among IQ). The distribution under the null hypothesis is empirically "generated" by permuting data labels.

It is important to note that permutation tests are not assumption free and do assume exchangeability (i.e., that when the null hypothesis is true the joint distribution of the observations remains unchanged under permutation of the group labels). Thus, we must be careful when permuting data labels. That is, we could reasonably assume $\varepsilon_{ij}$ and $\varepsilon_{i'j'}$ are independent and from the same distribution if individuals $i$, $j$, $i'$ and $j'$ are all unique. However, if there is overlap between individuals ($i$, $j$, $i'$ and $j'$ are *not* all unique), we should not expect this assumption to hold.

To illustrate how to randomly permute across individuals rather than all observations, consider a situation where we want to permute the residuals, $\mathrm{r}_{ij}$. If we were permuting across all observations, we would randomly shuffle the $\mathrm{r}_{ij}$ column. To permutate across individuals (therefore preserving exchangeability), we instead randomly shuffle the individual labels. See Figure S1 for an example:

| $\boldsymbol{i}$ | $\boldsymbol{j}$ | $\mathbf{r}_{ij}$ | → | $\boldsymbol{i}$ | $\boldsymbol{j}$ | $\mathbf{r}_{ij}$ |
|---|---|---|---|---|---|---|
| Alpha | Bravo | 1 | | Charlie | Alpha | 1 |
| Alpha | Charlie | 2 | **sample permutation:** | Charlie | Delta | 2 |
| Alpha | Delta | 3 | Alpha -> Charlie | Charlie | Bravo | 3 |
| Bravo | Charlie | 1 | Bravo -> Alpha | Alpha | Delta | 1 |
| Bravo | Delta | 2 | Charlie -> Delta | Alpha | Bravo | 2 |
| Charlie | Delta | 1 | Delta -> Bravo | Delta | Bravo | 1 |

*Figure S1: To illustrate how to randomly permute across individuals rather than all observations, consider a situation where we want to permute the residuals, $\mathrm{r}_{ij}$. If we were permuting across all observations, we would randomly shuffle the $\mathrm{r}_{ij}$ column. To permutate across individuals (therefore preserving exchangeability), we instead randomly shuffle the individual labels.*

We employed the Freedman-Lane approach (Freedman and Lane, 1983), while permuting across individuals to preserve exchangeability. The following is a (somewhat modified) explanation of the Freedman-Lane approach found on the package vignette for `permuco` on CRAN (Frossard and Renaud, 2019b) :

> The default method of `permuco` is the freedman_lane method that works as follows: we first fit the "small" model which only uses the nuisance variables $\boldsymbol{X_{con}}$ as predictors. Then, we permute its residuals and add them to the fitted values. These steps produce the permuted response variable $\boldsymbol{Dist}^*$ which constitutes the "new sample". It is fitted using the unchanged design $\boldsymbol{X_{con}}$ and $\boldsymbol{X_{coi}}$. In this procedure, only the residuals are permuted and they are supposed to share the same expectation

(of zero) under the null hypothesis. For each permutation, the effect of nuisance variables is hence reduced. Using the above notation, the fitted values of the "small" model can be written as $\boldsymbol{H}_{\boldsymbol{X}_{con}}\boldsymbol{Dist}$ and its residuals $\boldsymbol{R}_{\boldsymbol{X}_{con}}\boldsymbol{Dist}$ (where for any design matrix $\boldsymbol{M}$, $\boldsymbol{H}_{\boldsymbol{M}} = \boldsymbol{M}(\boldsymbol{M}^{\boldsymbol{T}}\boldsymbol{M})^{-\boldsymbol{1}}\boldsymbol{M}^{\boldsymbol{T}}$ is the "hat" matrix and $\boldsymbol{R}_{\boldsymbol{M}} = \boldsymbol{I} - \boldsymbol{M}(\boldsymbol{M}^{\boldsymbol{T}}\boldsymbol{M})^{-\boldsymbol{1}}\boldsymbol{M}^{\boldsymbol{T}}$ is the "residual" matrix). Its permuted version is pre-multiplied by a permutation matrix, e.g., $\boldsymbol{P}\boldsymbol{R}_{\boldsymbol{X}_{con}}\boldsymbol{Dist}$ . The permuted response variable is therefore simply written as $\boldsymbol{Dist}^{*} = \boldsymbol{H}_{\boldsymbol{X}_{con}}\boldsymbol{Dist} + \boldsymbol{P}\boldsymbol{R}_{\boldsymbol{X}_{con}}\boldsymbol{Dist} = (\boldsymbol{H}_{\boldsymbol{X}_{con}} + \boldsymbol{P}\boldsymbol{R}_{\boldsymbol{X}_{con}})\boldsymbol{Dist}$. The permuted F statistics are then computed using $\boldsymbol{Dist}^{*}$ and the unchanged design matrices $\boldsymbol{X}^{*}_{\boldsymbol{con}}$ = $\boldsymbol{X}_{\boldsymbol{con}}$ and $\boldsymbol{X}^{*}_{\boldsymbol{coi}}$ = $\boldsymbol{X}_{\boldsymbol{coi}}$.

#### 3.2. Results

##### *3.2.4. Canberra and Minkowski*

All remaining distance metrics are encapsulated in this section: Canberra and Minkowski of order 0.5, 1.4, 2 (Euclidean), and 3. All methods (not including F-test) adequately controlled type I error. Simulations 1-3, IQ: The GLS, F-test with ILE, and Mixed Model methods reached the power threshold within 20% signal. The permutation method reached the minimum power requirement within 40-60% of signal. Simulations 1-3, Treatment: All methods reached the power threshold within 20% signal. Simulation 4, IQ: The GLS, F-test with ILE, and Mixed Model methods reached the power threshold within 50% signal. The permutation method reached the minimum power requirement within 80-90% of signal. Simulation 4, Treatment: All methods reached the power threshold within 40-50% signal.

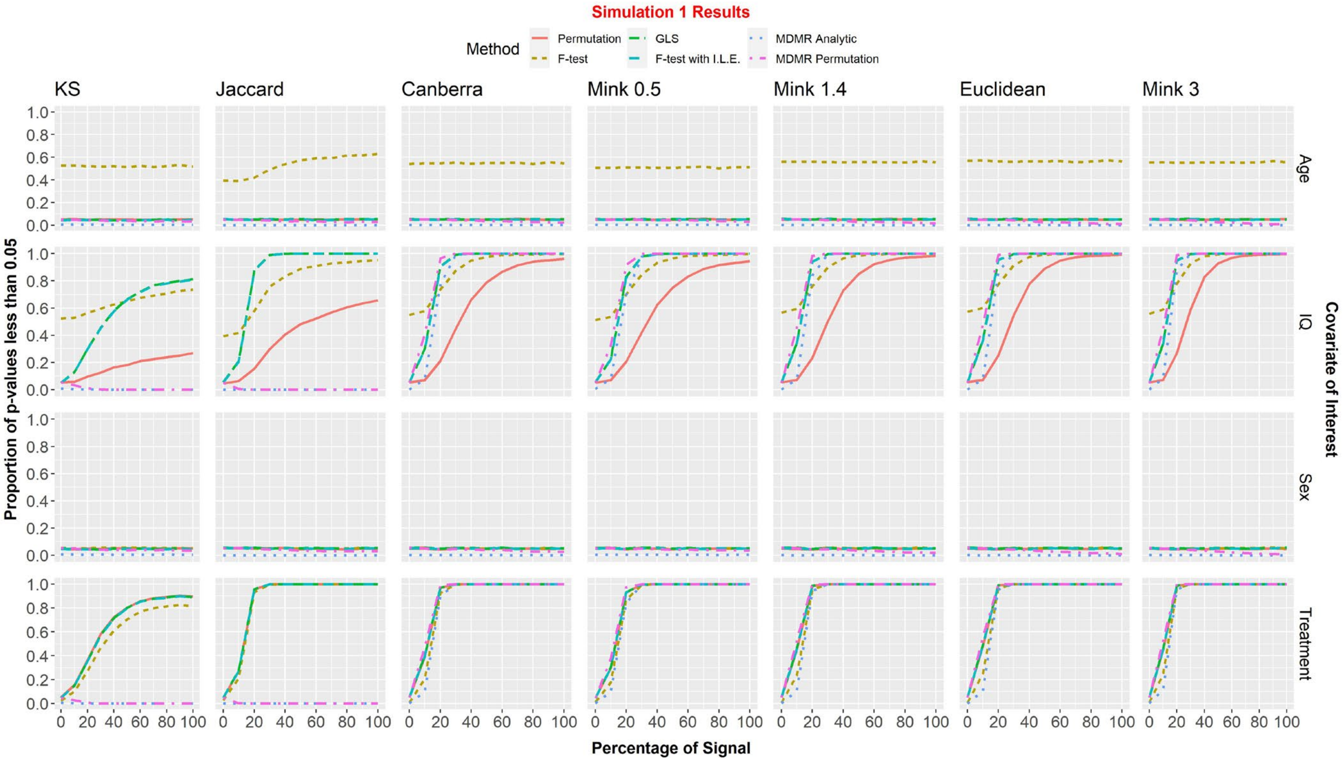


*Figure S2: Simulation 1 Results*

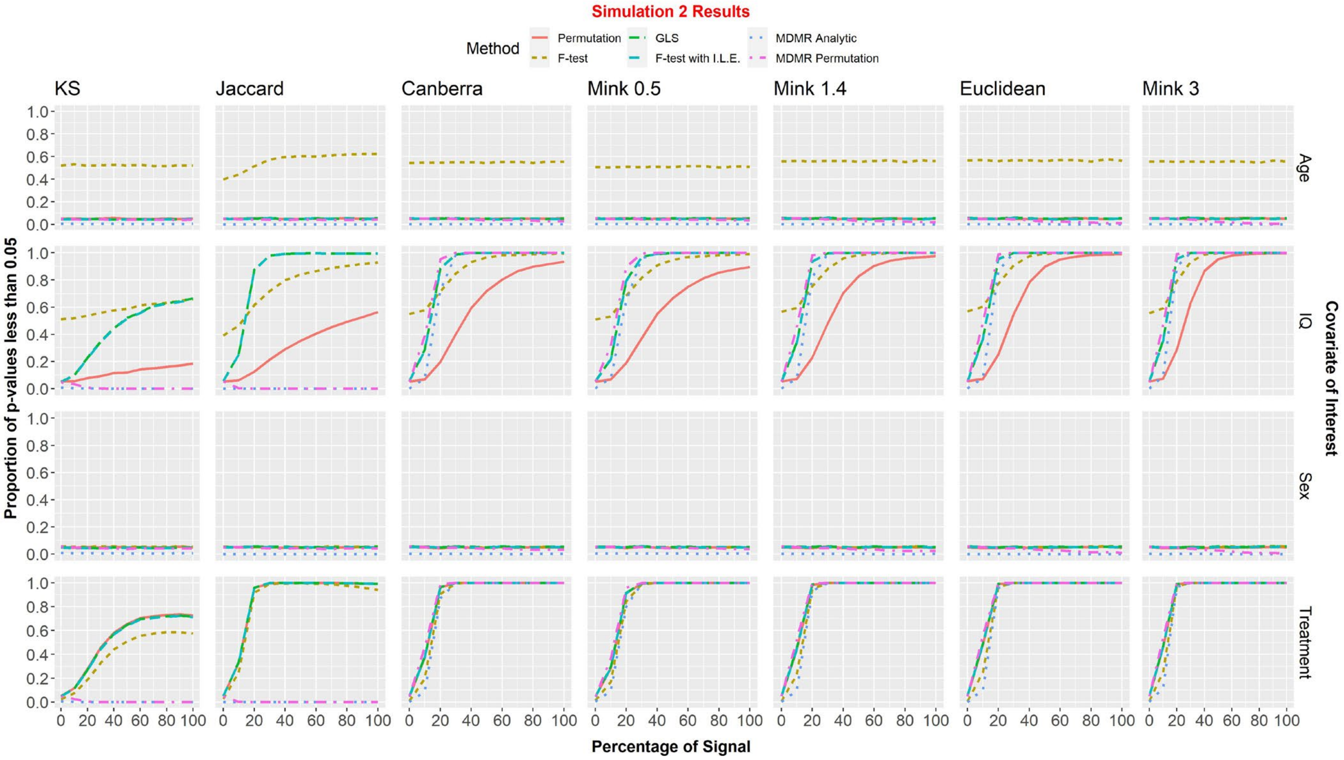


*Figure S3: Simulation 2 Results*

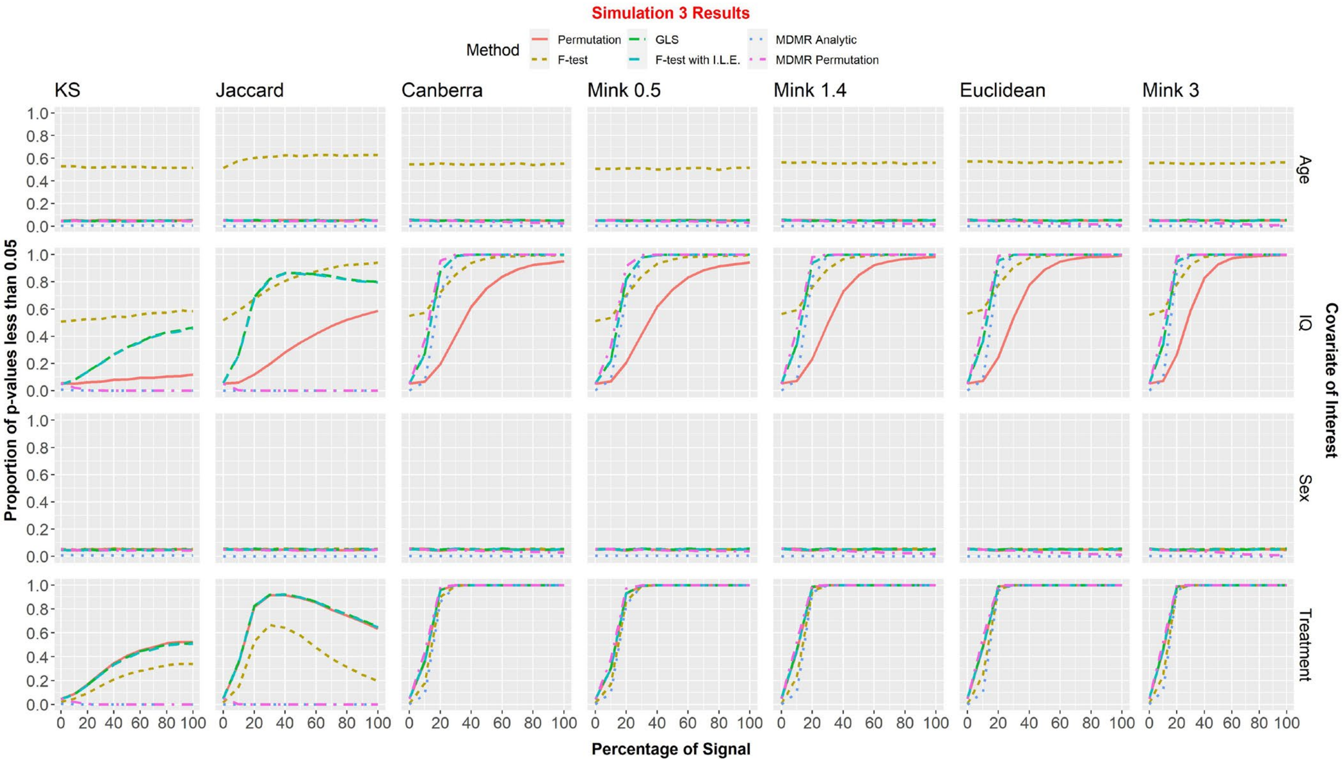


*Figure S4: Simulation 3 Results*

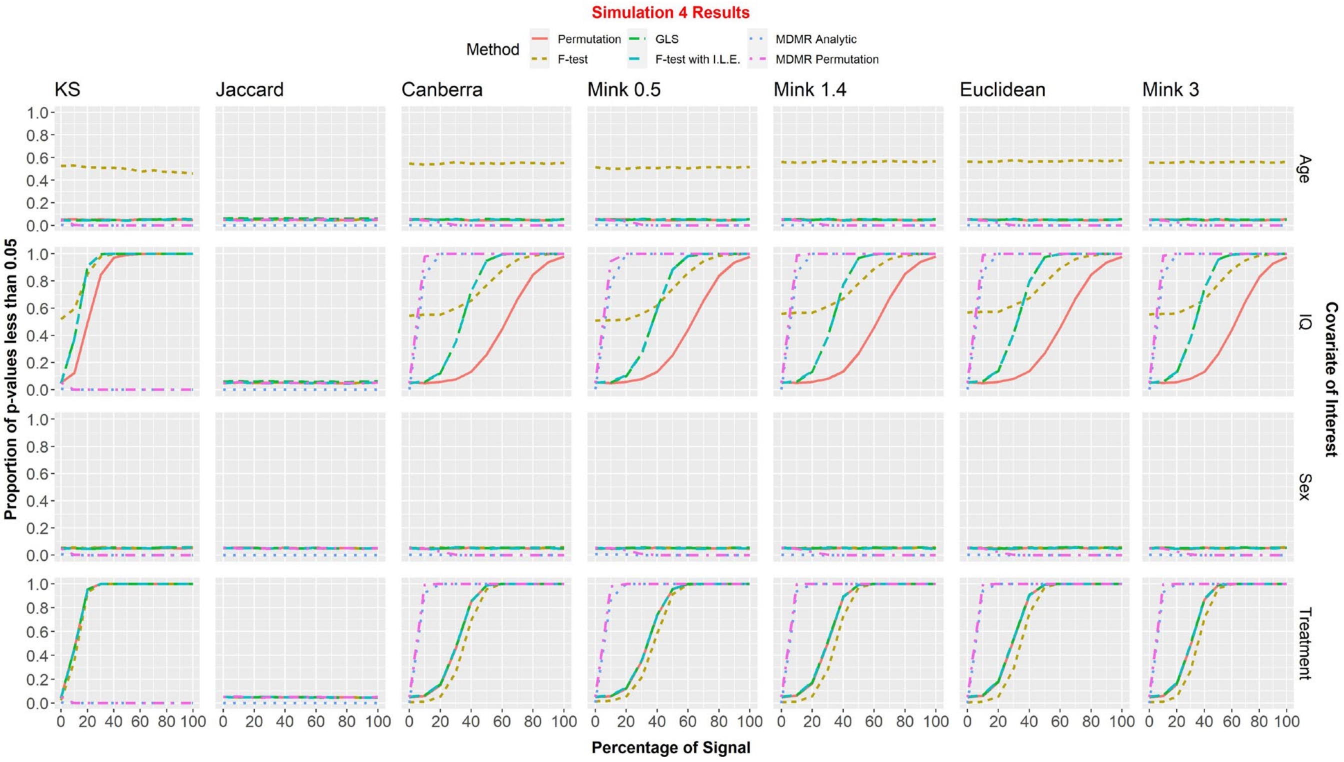


*Figure S5: Simulation 4 Results*

#### Appendix. Supplementary materials

Tables S1 and S2 contain the parameter estimates and standard errors from the HCP analysis described in the main text of the paper.

| | Resting State | | | | | | | |
|---|---|---|---|---|---|---|---|---|
| | KS | | Jaccard | | Mink 1.4 | | Euclidean | |
| | Est. | Std. Err. | Est. | Std. Err. | Est. | Std. Err. | Est. | Std. Err. |
| Fluid Intelligence | 2.51E-03 | 4.36E-04 | -9.72E-05 | 3.53E-05 | 2.83E-01 | 4.51E-02 | 9.09E-02 | 1.49E-02 |
| Age | 1.54E-03 | 5.41E-04 | -8.13E-05 | 4.38E-05 | 2.38E-01 | 5.59E-02 | 7.24E-02 | 1.85E-02 |
| Alcohol Abuse | 2.36E-02 | 6.00E-03 | 3.79E-04 | 4.86E-04 | 1.44E+00 | 6.20E-01 | 5.22E-01 | 2.05E-01 |
| Alc. Dependence | -1.12E-02 | 1.08E-02 | 1.06E-03 | 8.77E-04 | -1.37E+00 | 1.12E+00 | -4.06E-01 | 3.71E-01 |
| BMI | 6.68E-04 | 4.37E-04 | -9.93E-05 | 3.53E-05 | 2.03E-01 | 4.51E-02 | 6.49E-02 | 1.49E-02 |
| Education | -1.62E-03 | 1.12E-03 | -9.22E-05 | 9.07E-05 | -1.16E-01 | 1.16E-01 | -3.19E-02 | 3.83E-02 |
| Ethnicity | -1.12E-03 | 7.43E-03 | -4.80E-04 | 6.02E-04 | 6.06E-01 | 7.68E-01 | 1.85E-01 | 2.54E-01 |
| Gender | 5.53E-03 | 2.81E-03 | -1.84E-03 | 2.28E-04 | 2.45E+00 | 2.90E-01 | 7.82E-01 | 9.61E-02 |
| Handedness | -3.79E-05 | 5.85E-05 | -1.07E-05 | 4.73E-06 | 9.14E-03 | 6.04E-03 | 3.17E-03 | 2.00E-03 |
| Income | -8.11E-04 | 9.25E-04 | -1.73E-04 | 7.49E-05 | 1.06E-01 | 9.55E-02 | 4.23E-02 | 3.16E-02 |
| Race | 1.41E-02 | 5.01E-03 | -1.50E-03 | 4.06E-04 | 3.92E+00 | 5.17E-01 | 1.24E+00 | 1.71E-01 |
| Smoking Status | 5.99E-03 | 4.91E-03 | -1.45E-04 | 3.98E-04 | 5.63E-01 | 5.08E-01 | 1.74E-01 | 1.68E-01 |
| Participant 1 | -1.13E+00 | 1.98E-02 | 4.04E-01 | 1.60E-03 | 1.36E+02 | 2.04E+00 | 4.67E+01 | 6.77E-01 |
| Participant 2 | -1.01E+00 | 1.98E-02 | 4.09E-01 | 1.60E-03 | 1.28E+02 | 2.05E+00 | 4.40E+01 | 6.77E-01 |
| ⋮ | ⋮ | ⋮ | ⋮ | ⋮ | ⋮ | ⋮ | ⋮ | ⋮ |
| Participant 397 | -9.66E-01 | 2.02E-02 | 3.59E-01 | 1.64E-03 | 1.43E+02 | 2.09E+00 | 4.86E+01 | 6.91E-01 |

*Table S1: Parameters and Standard Errors for HCP resting state brain scans (nodal degree vectors) when modeled with our given regression framework and tested using the standard F-test with fixed individual level effects.*

| | Working Memory | | | | | | | |
|---|---|---|---|---|---|---|---|---|
| | KS | | Jaccard | | Mink 1.4 | | Euclidean | |
| | Est. | Std. Err. | Est. | Std. Err. | Est. | Std. Err. | Est. | Std. Err. |
| Fluid Intelligence | 5.33E-04 | 4.57E-04 | -5.93E-05 | 2.81E-05 | 7.08E-02 | 3.60E-02 | 2.69E-02 | 1.22E-02 |
| Age | 9.69E-04 | 5.67E-04 | -1.15E-04 | 3.48E-05 | 2.59E-01 | 4.46E-02 | 8.72E-02 | 1.52E-02 |
| Alcohol Abuse | -1.77E-03 | 6.29E-03 | 4.92E-04 | 3.86E-04 | -4.12E-01 | 4.94E-01 | -1.29E-01 | 1.68E-01 |
| Alc. Dependence | 3.10E-03 | 1.14E-02 | 1.23E-03 | 6.97E-04 | -1.27E+00 | 8.93E-01 | -4.02E-01 | 3.04E-01 |
| BMI | 1.12E-03 | 4.57E-04 | 3.93E-05 | 2.81E-05 | 2.40E-02 | 3.60E-02 | 1.04E-02 | 1.23E-02 |
| Education | 3.06E-03 | 1.17E-03 | -2.61E-04 | 7.21E-05 | 2.16E-01 | 9.23E-02 | 7.30E-02 | 3.15E-02 |
| Ethnicity | 2.84E-03 | 7.79E-03 | 6.98E-04 | 4.78E-04 | -5.22E-01 | 6.12E-01 | -1.60E-01 | 2.09E-01 |
| Gender | 1.04E-03 | 2.95E-03 | -1.11E-03 | 1.81E-04 | 1.30E+00 | 2.32E-01 | 4.52E-01 | 7.89E-02 |
| Handedness | -5.50E-05 | 6.13E-05 | -4.51E-06 | 3.76E-06 | 1.58E-03 | 4.82E-03 | 6.00E-04 | 1.64E-03 |
| Income | 3.62E-04 | 9.69E-04 | -3.12E-05 | 5.95E-05 | 6.58E-02 | 7.62E-02 | 2.33E-02 | 2.60E-02 |
| Race | 5.35E-03 | 5.25E-03 | -1.70E-03 | 3.22E-04 | 1.21E+00 | 4.13E-01 | 3.86E-01 | 1.41E-01 |
| Smoking Status | 2.89E-03 | 5.15E-03 | 1.49E-04 | 3.16E-04 | 5.12E-02 | 4.05E-01 | -2.39E-02 | 1.38E-01 |
| Participant 1 | -9.75E-01 | 2.07E-02 | 3.74E-01 | 1.27E-03 | 1.57E+02 | 1.63E+00 | 5.34E+01 | 5.55E-01 |
| Participant 2 | -9.98E-01 | 2.08E-02 | 3.69E-01 | 1.27E-03 | 2.03E+02 | 1.63E+00 | 7.01E+01 | 5.56E-01 |

| | | | | | | | | |
|---|---|---|---|---|---|---|---|---|
| ⋮ | ⋮ | ⋮ | ⋮ | ⋮ | ⋮ | ⋮ | ⋮ | ⋮ |
| Participant 397 | -3.14E-01 | 2.12E-02 | 3.78E-01 | 1.30E-03 | 4.14E+02 | 1.67E+00 | 1.43E+02 | 5.67E-01 |

*Table S2: Parameters and Standard Errors for HCP working memory brain scans (nodal degree vectors) when modeled with our given regression framework and tested using the standard F-test with fixed individual level effects.*

In addition to our primary analysis detailed in the main text, we also examined these relationships with respect to differences in modularity between individuals employing Scaled Inclusivity (SI) given that modularity analyses can capture the spatial distribution of intrinsic brain networks that are associated with various cognitive tasks (Moussa et al., 2012). Scaled Inclusivity is a nodal measure of spatial consistency in modular structure when compared across study participants or to a predefined Region of Interest (ROI). The SI values in any given voxel indicate how well the module that includes that voxel overlaps with the module for that voxel in other participants or with the chose ROI. Details on the computation of SI can be found in prior work (Moussa et al., 2012; Steen et al., 2011). We compare each subject's modular structure to an ROI encompassing the Default Mode Network (DMN) or the Central Executive Network (CEN). Resting state data were compared to the DMN and Task data were compared to the CEN. The different ROIs were selected because they represent the expected subnetworks based on the type of scan. For each participant, whole brain SI images were generated with the value in each voxel being the SI value computed for that voxel using the specific ROI. Whole-brain SI maps were averaged across participants to generate group maps shown in Figure S1.

The whole-brain SI images from each participant were transformed into weighted Scaled Inclusivity and the analysis follows the same process as before (with degree vectors): Key nodes of interest (binary vectors used for the Jaccard index) based on nodal SI were identified, selecting the top 20% highest SI nodes. KS statistic, Minkowski distance of order 1.4, and Euclidean distance (Minkowski of order 2) were calculated for each pair of individuals using their nodal SI vectors. The Jaccard distance was calculated for each pair of individuals using their binary SI vectors.

Distance covariates for each pair of individuals were calculated. A continuous variable's distance (Age, for instance) was calculated as $|Age_i - Age_j|$ for the pair of individuals $i$ and $j$. A binary or categorical variable's distance (Education, for instance) was calculated as $\mathbb{1}\{Edu_i \neq Edu_j\}$ for the pair of individuals $i$ and $j$.

We evaluated differences between networks using the Standard F-test with Individual Level Fixed Effects. A complete list of p-values for both resting state (SI calculated with Default Mode Network, DMN) and working memory (SI calculated with Central Executive Network, CEN) can be seen in Table S3.

| | Resting State (DMN Scaled Inclusivity) | | | | Working Memory (CEN Scaled Inclusivity) | | | |
|---|---|---|---|---|---|---|---|---|
| | KS | Jaccard | Mink 1.4 | EUC | KS | Jaccard | Mink 1.4 | EUC |
| Fluid Intelligence | 5.32E-01 | 5.24E-02 | 9.69E-03 | 8.59E-03 | 1.30E-03 | 3.86E-03 | 6.07E-02 | 1.52E-01 |
| Age | 2.80E-01 | 5.90E-01 | 2.22E-03 | 7.55E-03 | 6.88E-01 | 9.82E-01 | 5.12E-01 | 5.38E-01 |
| Alcohol Abuse | 3.70E-01 | 4.32E-01 | 2.89E-01 | 3.87E-01 | 3.12E-01 | 6.42E-01 | 4.46E-01 | 4.45E-01 |
| Alc. Dependence | 9.63E-01 | 6.80E-01 | 3.26E-03 | 4.75E-04 | 4.01E-01 | 3.95E-01 | 2.74E-01 | 3.21E-01 |
| BMI | 2.24E-03 | 9.36E-01 | 9.74E-01 | 8.23E-01 | 8.79E-01 | 1.52E-01 | 9.07E-01 | 8.36E-01 |
| Education | 2.15E-01 | 6.60E-01 | 7.39E-01 | 7.57E-01 | 5.60E-01 | 2.68E-01 | 9.83E-01 | 9.58E-01 |
| Ethnicity | 3.71E-02 | 2.80E-01 | 5.83E-01 | 4.31E-01 | 5.61E-01 | 1.10E-01 | 4.96E-01 | 4.93E-01 |
| Gender | 1.62E-01 | 7.67E-11 | 1.19E-12 | 8.37E-12 | 6.53E-01 | 4.79E-08 | 1.42E-03 | 9.57E-03 |
| Handedness | 3.90E-01 | 6.75E-01 | 4.07E-02 | 1.28E-01 | 1.86E-01 | 2.90E-01 | 7.11E-01 | 8.85E-01 |
| Income | 5.80E-01 | 6.37E-01 | 5.23E-01 | 5.17E-01 | 7.25E-02 | 7.08E-01 | 4.69E-01 | 3.45E-01 |
| Race | 6.96E-01 | 1.06E-03 | 4.19E-03 | 4.06E-03 | 8.15E-01 | 3.54E-10 | 6.90E-04 | 1.60E-03 |
| Smoking Status | 8.90E-01 | 4.66E-01 | 7.83E-01 | 6.89E-01 | 9.26E-01 | 3.24E-01 | 2.49E-01 | 2.81E-01 |

Legend **0** **0.05** **1**

*Table S3: P-values for HCP scaled inclusivity resting state and working memory brain scans when modeled with our given regression framework and tested using the standard F-test with fixed individual level effects.*

For maps showing community organization of the DMN and CEN networks at rest and during a working memory task, see Figure S9.

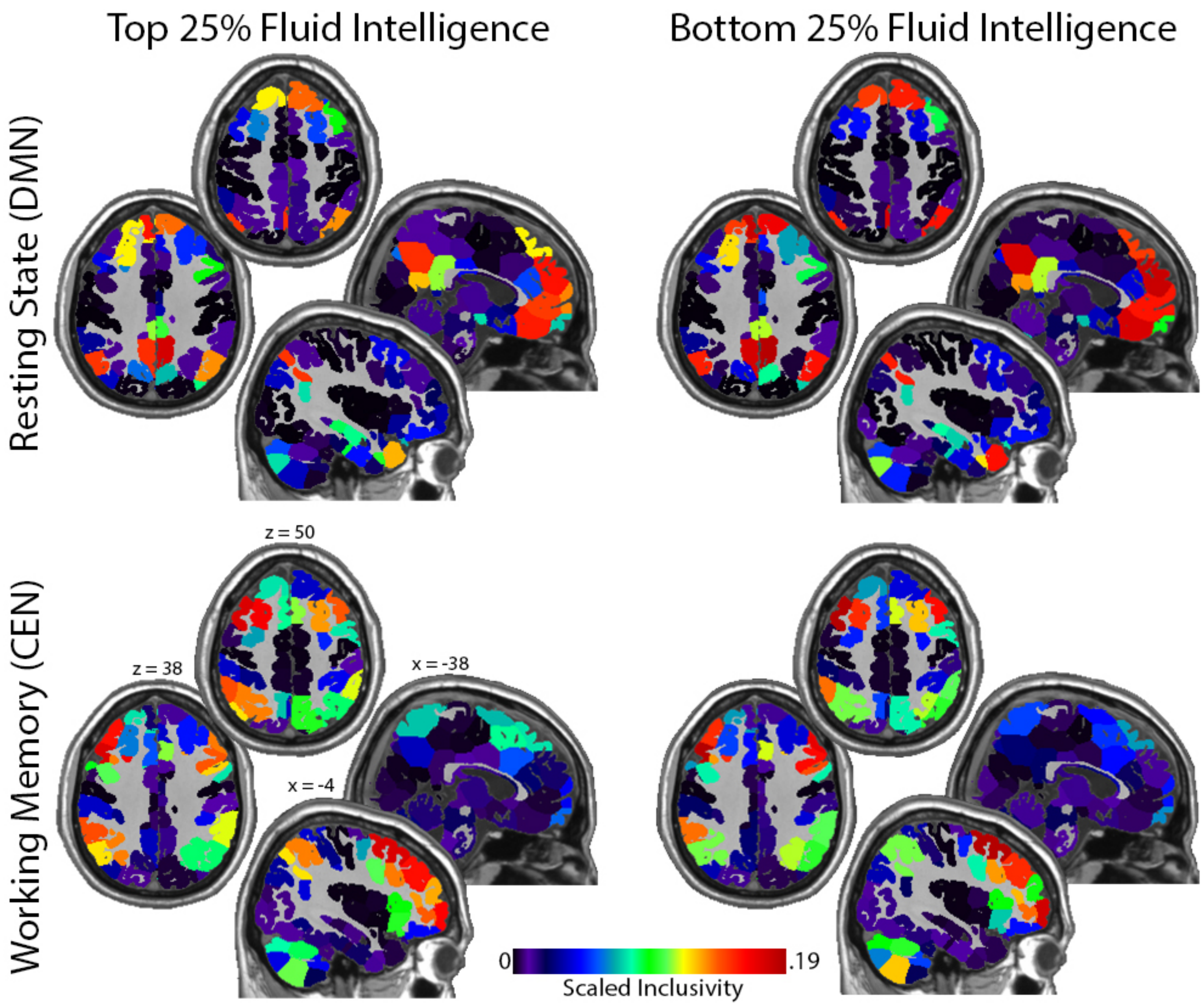


*Figure S6: Maps showing community organization of the DMN and CEN networks at rest and during a working memory task, respectively. Maps are shown for the top and bottom intelligence quartiles. The scaled inclusivity (SI) values indicate the likelihood that a region was including in the community associated with each intrinsic brain network. The maps represent a "average" community in each group. The significant association between SI and intelligence is demonstrated by increases in community structure for both networks with decreases in fluid intelligence. The association is most prominent in the DMN but can is also evident in the frontal aspects of the CEN, particularly on the right. Each quadrant shows 2 axial and 2 sagittal images. The Montreal neurological Institute (MNI) coordinates shown in the bottom left quadrant apply to all quadrants. Calibration bar applies to all images.*
